# Pooled screening of CAR T cells identifies non-native signaling domains for next-generation immunotherapies

**DOI:** 10.1101/2021.07.11.451980

**Authors:** Daniel B. Goodman, Camillia S. Azimi, Kendall Kearns, Kiavash Garakani, Julie Garcia, Nisarg Patel, Byungjin Hwang, David Lee, Emily Park, Chun Jimmie Ye, Alex Marson, Jeff A. Bluestone, Kole T. Roybal

## Abstract

Chimeric antigen receptors (CARs) repurpose natural signaling components to retarget T cells to refractory cancers, but have shown limited efficacy against solid tumors. Here, we introduce ‘CAR Pooling’, a multiplexed approach to rapidly identify CAR designs with clinical potential. Forty CARs with diverse immune costimulatory domains were assessed in pooled assays for their ability to stimulate critical T cell effector functions during repetitive stimulation that mimics long-term tumor antigen exposure. Several non-native domains from the TNF receptor family exhibited enhanced proliferation (CD40) or cytotoxicity (BAFF-R and TACI) relative to clinical benchmarks, and fell into distinct states of memory, cytotoxicity, and metabolism. BAFF-R CAR T cells were enriched for a highly cytotoxic and NK-cell-like innate phenotype previously associated with positive clinical outcomes. ‘CAR Pooling’ enables efficient exploration of how CAR design affects cell activity and can be applied to optimize receptors across a range of applications and cell types.

## INTRODUCTION

Adoptive cell therapy using engineered chimeric antigen receptor (CAR) T cells has revolutionized the treatment of B-cell leukemias and lymphomas (Ahmad et al., 2020; Azimi et al., 2019). CARs currently in the clinic use either a 4-1BB or CD28 intracellular costimulatory domain, both of which come from natural, well-studied T cell costimulatory receptors. Costimulation is a critical component of immune activation, and CARs lacking the ‘signal 2’ from a costimulatory domain quickly become anergic upon stimulation(Chen and Flies, 2013). CARs containing 4-1BB and CD28 intracellular domains are used in second generation CARs, which elicit more robust and sustained T cell activation than the original CD3ζ-only CARs. While both CD28- and 41BB-containing CARs are effective therapeutics, there are substantial differences in their synapse development, cytotoxicity, metabolic state, and clinical performance (Davenport et al., 2018; Kawalekar et al., 2016; Weinkove et al., 2019; Zhao et al., 2020). The current state of the literature indicates that T cells expressing the CD28 CAR are initially faster to proliferate and kill tumor cells but suffer from reduced long-term engraftment and heightened exhaustion after prolonged activation (Maude et al., 2014, 2018; Park et al., 2018; Salter et al., 2018; Ying et al., 2019).

There is considerable diversity in costimulatory domains, and evidence for both quantitative and qualitative differences in costimulatory signaling in the context of a CAR (Weinkove et al., 2019). 4-1BB and CD28 utilize two separate signaling pathways (TRAF and PI3K/Lck), however these pathways converge upon conserved signaling intermediates, suggesting that costimulatory domains from other immune cells may also be able to signal in T cells. Other studies have characterized additional T cell costimulatory domains individually within CARs or searched for mutant domains with enhanced properties(Dawson et al., 2020; Guedan et al., 2014, 2018, 2020; Kagoya et al., 2018). However, the scale of these searches has been limited to selected domains, focused primarily on receptors with known functions in T cells.

Pooled screens have been a powerful tool for probing T cell biology, including using CRISPR knockouts and switch receptors (Roth et al., 2020; Shifrut et al.), but have yet to be applied to CAR engineering. Pooled assays offer both increased throughput and direct comparison of cells from the same blood donor tested in exactly the same conditions. Screening of large numbers of domains to assess their effects on multiple cell-intrinsic T cell phenotypes would help identify optimal CAR designs for clinical applications. While pooled measurement can be applied to many aspects of CAR architecture, signaling domains, which are small, have minimal secondary structure, and consist of short and modular signaling motifs (Diella et al., 2008), lend themselves to pooled characterization using large synthetic DNA libraries (Kosuri and Church, 2014). Whole domains, individual motifs, and broad design principles identified from the data can then be leveraged to generate focused sets of CARs for in-depth *in vitro* and *in vivo* analysis and the potential for clinical translation.

The lack of persistence and long-term efficacy in patients is a central problem for current CAR T therapies (Kershaw et al., 2006; Park et al., 2007). To mimic the protracted stress of chronic antigen exposure in difficult-to-eliminate solid tumors, we developed a four-week *in vitro* repetitive stimulation assay and measured cytokine production, proliferation, and persistence to identify costimulatory domains that improved the long-term anti-tumor activity of CAR T cells. We assembled a cosignaling library – consisting of both inhibitory and stimulatory domains – from a range of innate and adaptive immune cells that utilize common signaling machinery present in T cells (Chen and Flies, 2013). We then performed a suite of pooled assays in primary human CD4 or CD8 T cells containing this CAR library, generating the first systematic survey of the CAR T cell costimulation landscape. We find that certain costimulatory domains have enhanced capabilities within either CD4 or CD8 T cells and that signaling domains normally associated with other immune lineages (macrophages, B cells, NK cells) are capable of strongly activating T cells. We identify a set of potent costimulatory domains from the TNF receptor family that lead to enhanced proliferation and cytotoxicity and have *in vivo* rejection dynamics equal to 4-1BB and CD28 in a solid tumor model. Additionally, we identify KLRG1, an inhibitory domain that silences activation and keeps the T cell in a naive transcriptional state. Finally, single-cell RNA and surface protein expression profiles among these candidates showed that CAR Ts using the signaling domain from BAFF-R are enriched for a highly cytotoxic expression signature. This signature, associated with IFN-γ production and an NK-cell-like innate phenotype, has been previously associated with enhanced 4-1BB CAR T engraftment and improved response against melanoma in clinical studies.

## RESULTS

### CAR Pooling: Generation and screening of a pooled library of CARs with a diverse suite of costimulatory domains

Innate and adaptive immune cells utilize accessory receptors to elicit critical cellular functions. These receptors often use modular linear signaling motifs and signaling proteins that are conserved across the immune system (Chen and Flies, 2013; Diella et al., 2008). We mined 40 costimulatory and coinhibitory receptor intracellular domains from different protein families and functional classes associated with several immune cell types, including NK cells, B cells, and innate immune cells (Figure 1A, 1B). To generate this ‘CAR Pooling’ library, we synthesized and cloned the domains into a second-generation CAR scaffold and, with T cells from two human donors, transduced a lentiviral library containing the entire pool of domains into separate CD4 and CD8 T cell cultures (Figure 1C). CAR positive T cells were sorted based on a T2A-GFP marker for a defined range of CAR expression and rested for 5 days before proceeding with tumor cell stimulation. Once generated, these novel CARs were studied for their effects on T cell proliferation and exhaustion after prolonged stimulation, in comparison to clinical benchmark CARs with 4-1BB or CD28 costimulatory domains.

**Figure 1:**
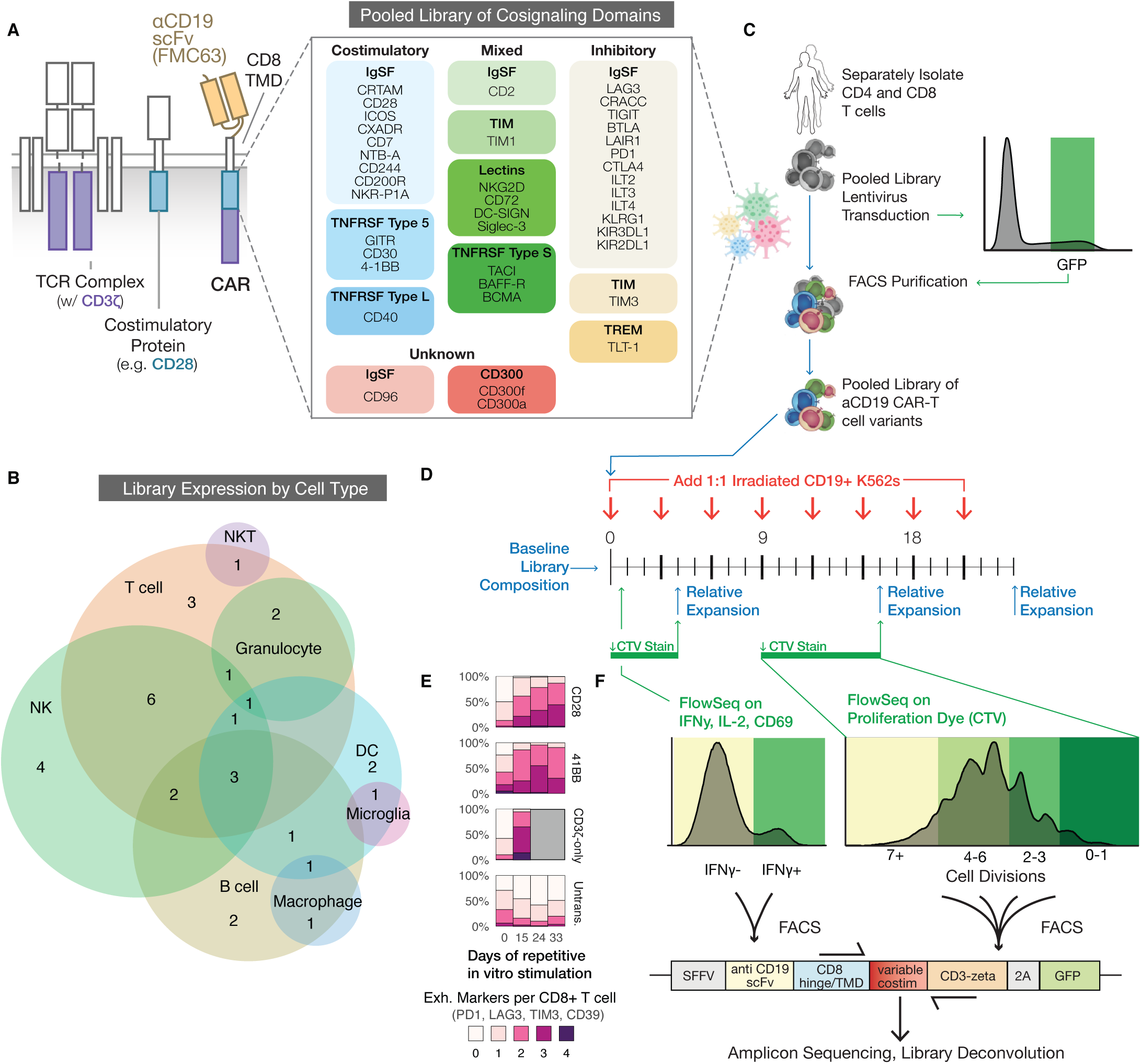
CAR Pooling: Generation and screening of a pooled library of CARs with a diverse suite of costimulatory domains. **A)** Our chimeric antigen receptors combine a ScFv against human CD19 (FMC63), a CD8 hinge and transmembrane domain (TMD), one of 40 different intracellular domains, and a CD3ζ domain (left). Cosignaling domains from across the human proteome were identified, codon-optimized, synthesized, pooled into a plasmid library, and packaged to generate lentivirus. **B)** A Venn diagram showing natural expression of library members across immune cell types. Expression patterns individually listed in (Table S1). **C)** Primary human CD4 and CD8 T cells were separately isolated from PBMCs for two human donors and lentivirally transduced with the library. The cells were subsequently purified via FACS using 2A-GFP fluorescence within one log of mean expression to reduce variability. **D)** The pooled library was repeatedly stimulated 1:1 with either CD19+ or CD19- K562 tumor cells to quantify antigen specific activation (CD69), CD4 cytokine production (IFN-γ, IL-2), and proliferation (cell trace violet--CTV). The library was also bulk-sequenced at the specified time points to measure relative expansion of individual constructs within the sample. **E)** Percentage of CD8+ T cells expressing different numbers of exhaustion markers (PD1, TIM3, LAG3, CD39) after a repeat stimulation assay with CD19+ irradiated K562 tumor cells. T cells expressing CD28, 4-1BB, or CD3ζ-only CARs are compared to untransduced cells. Grey boxes correspond to timepoints in which no live cells remained in culture. See Figure S1A/B for a second donor and data for CD4+ T cells. **F)** We used FlowSeq, a FACS- and NGS-based pooled quantification workflow, to quantify enrichment by sorting the library into bins of fluorescent signal corresponding to a functional readout, as shown by the colored boxes on the representative histograms. We then directly amplified the costimulatory domain via genomic DNA extraction and PCR, and performed NGS amplicon sequencing on each bin to estimate the phenotype for each library member.

To mimic the exhaustion conditions encountered in patients with a high tumor burden, we performed repetitive, long-term stimulations of the CAR T cells over 24-33 days. This was accomplished via an *in vitro* rechallenge model where T cells were stimulated with additional CD19+ or CD19- K562 cells, at a constant 1:1 T cell to target ratio, every three days (Figure 1D). K562 cells were irradiated prior to their addition to reduce their proliferative capacity and prevent rapid depletion of the media. After testing CD28 and 4-1BB CAR T cells in this assay over 33 days of repeated stimulations, the T cells became increasingly exhausted, based on measurement of four exhaustion markers (PD1, TIM3, LAG3, CD39) (Figure 1E, S1A, S1B). Additionally, a CAR containing only the CD3ζ signaling domain expressed a significant exhaustion phenotype by day 15 and did not survive after day 24 of culture, indicating that the effects of costimulation, which are critical to durable responses *in vivo*, are at least partially captured by this *in vitro* model (Esensten et al., 2016; Finney et al., 1998; Lafferty and Cunningham, 1975; Weinkove et al., 2019).

To efficiently characterize the ‘CAR Pooling’ library across multiple assays and time points, we employed FlowSeq, a pooled measurement approach that combines FACS and amplicon sequencing to quantitatively measure any fluorescence-based readout across a genetically-diverse population of cells (Goodman et al., 2013). We used FlowSeq to measure different markers of T cell function, such as activation (CD69), cytokine production (IFN-γ and IL-2), and proliferation using Cell Trace Violet (CTV) dye. For each pooled assay, cells were sorted into either two bins (for marker expression) or four bins (for CTV dye dilution) and separately sequenced to compare the functional differences between the domains for each assay ( Figure 1D, 1F). This multiplexed screening approach allowed us to compare the extent to which domains differentially affect CAR T activity between CD4 and CD8 T cell types and between early and late stages of antigen stimulation and expansion.

### Multidimensional comparison of cosignaling domains across multiple weeks of expansion identifies a subset with potent costimulatory activity

Antigen-induced proliferation varied across CAR T cells expressing different costimulatory domains (Figure 2A). Despite donor variability, the relative magnitude of proliferation across replicates was consistent (Figure S1C), with the canonical CD28 and 4-1BB costimulatory domains often among the domains that promoted the most proliferation (Figure 2A). We also measured IL-2 and IFN-γ secretion in CD4 CAR T cells (Figure 2B) and CD69 expression in both CD4 and CD8 T cells (Figure S1D). CD28 CD4 CAR T cells expressed high levels of cytokines, producing more IL-2 than any other domain. While some cytokine secretion is desired, high levels of cytokine production has been previously associated with CD28 CAR T activation-induced cell death (AICD) (Künkele et al., 2015), T cell exhaustion (Beltra et al., 2014; Liu et al., 2021) and higher rates of cytokine release syndrome (CRS) within patients (Zhao et al., 2020). This suggests that overproduction of cytokines may be less ideal for clinical use. Notably, 4-1BB, BAFF-R, TACI, and NTB-A showed more moderate but enhanced cytokine production compared to the population average.

**Figure 2:**
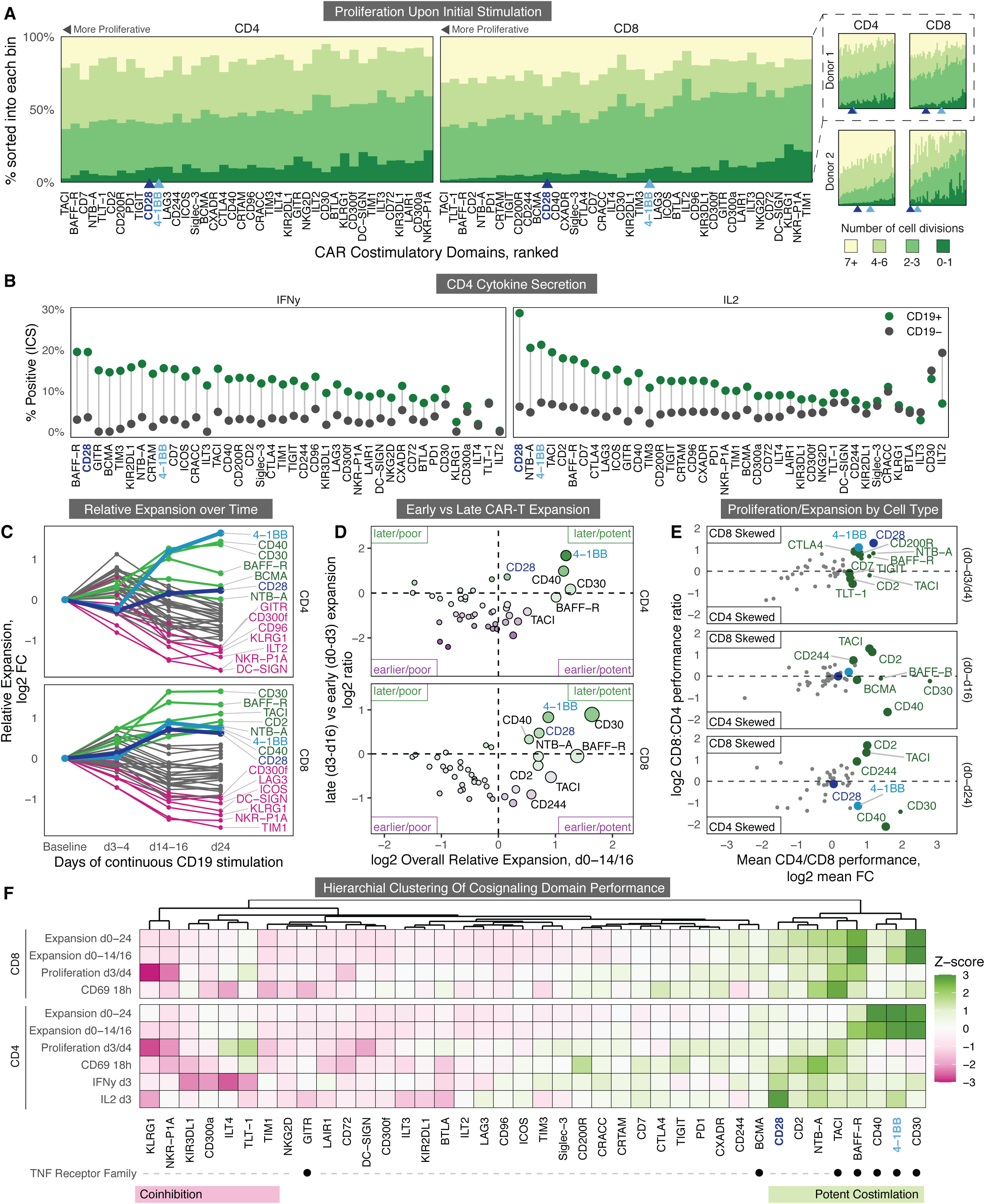
Multidimensional comparison of cosignaling domains across multiple weeks of expansion identifies a subset with potent costimulatory activity. **A)** FlowSeq measurement of proliferation of CD4 (left) and CD8 (right) T cells containing different CAR library domains, separately stimulated *in vitro* with irradiated CAR Ts for 3 or 4 days (see Figure 1C). Percentages of cells with different numbers of divisions were calculated from amplicon sequencing of sorted CTV bins. CARs are ranked from left to right by the average number of cell divisions in CD4 or CD8 cells from highest (left) to lowest (right). CD28 and 4-1BB CARs are highlighted in blue shades. On the right, two donors are shown separately along with the rankings of CD28 and 4-1BB. **B)** FlowSeq measurement of intracellular cytokine accumulation across library domains in CD4 cells, 18 hours after the initial addition of either CD19+ or CD19- irradiated K562s. Cells are ranked based on the difference in percent cytokine-positive cells between CD19+ and CD19- conditions. **C)** Relative expansion over time of the CAR domain library in CD4 and CD8 T cells, based on the average fold-change in library abundance from the initial library before stimulation. Mean of 3 replicates and two donors. The 6 CAR domains with the most and least relative expansion are labelled in green and pink, respectively, with CD28 and 4-1BB labelled in blue shades. **D)** Comparison of early vs late antigen-stimulated proliferation. The X axis measures overall expansion by day 14 or 16 (d14/16) with more potent CARs on the right and less potent CARs on the left. The Y axis measures the ratio of late proliferation (d9-d16) vs early proliferation (d0-d3). CARs above 0 on the Y axis are more expanded in the library at later time points, and CARs below 0 are more expanded earlier. Top right quadrant indicates that the most potent proliferators overall saw relatively less early expansion in the first three days, while the less potent CARs are in the bottom quadrants, showing more early expansion. **E)** Comparison of CD4 versus CD8 proliferation and expansion after CD19+ K562 stimulation. CARs in green have statistically significant increased expansion (DESeq2) in the assay in either CD4 or CD8 T cells. Larger green dots correspond to constructs that have significantly better performance in either CD4s or CD8s. The X axis is the mean fold-change across CD4 and CD8 T cells, while the Y axis is the ratio between CD8 and CD4 expansion. **F)** Hierarchical clustering of CD19+ assays across CAR library domains in CD4 and CD8 T cells. All 40 domains were clustered based on their z-scores in each assay, identifying a distinct cluster of 8 potent costimulatory domains shown on the right (green bar). A small set of domains which perform poorly in various aspects of CAR function are on the left (pink bar). TNF receptor family members are noted with black dots.

The lack of long-term CAR T cell persistence is often cited as a major reason for antigen-positive relapse in patients (Jafarzadeh et al., 2020). Measuring changes in library composition over time integrates multiple aspects of CAR T cell dynamics that underlie persistence, including proliferation and cell lifespan. Amplicon sequencing of the library immediately before stimulation allowed for an assessment of the baseline library distribution, and the library composition was subsequently measured after the first, sixth, and eighth subsequent stimulations (days 3 or 4, 14 or 16, and 24 respectively). Using these measurements, we were able to track the relative expansion and contraction of each CAR in the library (Figure 2C). Many of the costimulatory domains which preferentially expanded after the first stimulation were diminished in the library after multiple stimulations, indicating that initial proliferation did not correlate well with long-term proliferative capacity and persistence (Figure 2D). We also saw differences between CD4 and CD8 expansion dynamics (Figure 2E). Most costimulatory domains saw a greater expansion overall in CD8 T cells, with the notable exceptions of the CD30, CD40 and 4-1BB CAR T cells, which, after 24 days of stimulation, expanded over 2-fold more in CD4s than in CD8s. As noted by previous studies, these results imply that using different combinations of costimulatory domains between CD4 and CD8 T cells may improve overall CAR T therapeutic efficacy (Guedan et al., 2018).

Antigen-independent proliferation also varied considerably among cells expressing the individual costimulatory domains. While overall proliferation was universally lower without antigen, strongly proliferative domains also exhibited enhanced proliferation in co-culture with CD19- K562 cells (Spearman’s ρ = 0.63-0.75, p = 1.3e^-5^), indicating that higher antigen-dependent proliferation was strongly correlated with increased basal proliferation (Figure S1F). Among the top-performing costimulatory domains, CD28 and TACI exhibited the highest degree of non-specific proliferation, while CD40 had the lowest (Figure S1F).

### Scoring CARs across pooled measurements identifies cosignaling domains with distinct stimulatory or inhibitory activity

To summarize the functionality of all domains in the library over the repetitive stimulations, hierarchical clustering was performed based on the relative performance of all domains (Figure 2F). A subset of CARs containing CD28, 4-1BB, and several additional domains clearly clustered into a potent costimulatory group, demonstrating enhanced T cell functions relative to the average costimulatory domain in the library (Figure 2F, right). A smaller set of domains clustered into a coinhibitory group with reduced proliferation and cytokine secretion across multiple assays (Figure 2F, left). Within the potent costimulatory group described above, hierarchical clustering further separated domains into three subsets. The first consisted of the 4-1BB, CD30, and CD40 domains, which demonstrated less initial proliferation and early cytokine secretion but were significantly enriched after multiple weeks of repetitive stimulation, particularly in CD4 T cells. The second subset of potent costimulatory domains – CD28, CD2, and NTB-A – showed stronger initial proliferation, CD69 expression, and cytokine secretion, but had less long-term expansion and persistence within the pool after repetitive stimulation. The final subset – BAFF-R and TACI – had an intermediate phenotype, with heightened initial proliferation in CD8s similar to CD30, moderate cytokine production, and enhanced CD69 expression in CD4s comparable to CD28. Overall, the six most potent CAR costimulatory domains were distributed along a spectrum between the strong early response of CD28 and enhanced late-stage expansion and CD4-bias of 4-1BB, suggesting that there may be inherent tradeoffs between these two aspects of CAR T activity.

To further compare the overall performance of the various costimulatory domains, a principal component analysis (PCA) was performed across all pooled measurements both with and without antigen stimulation (Figure 3A, Figure S2A-C). The PCA showed that the individual domains are spread across a diverse costimulatory landscape, with PC1 being associated with early proliferation, cytokine secretion, CD69 activation, and tonic signaling, while PC2 is associated with long-term expansion in CD8s, less cytokine secretion and early proliferation, and relative lack of tonic signaling (Figure S2A). PC2 also showed a significant correlation with the size of the costimulatory domain, suggesting that increasing the distance between the membrane and CD3ζ reduces strength of early activation and tonic signaling (Figure S2C). This is supported by recent work showing that altering the position of individual ITAM motifs within CD3ζ can modulate differentiation and memory formation (Feucht et al., 2019; Holst et al., 2008; James, 2018; Xu et al., 2008).

**Figure 3:**
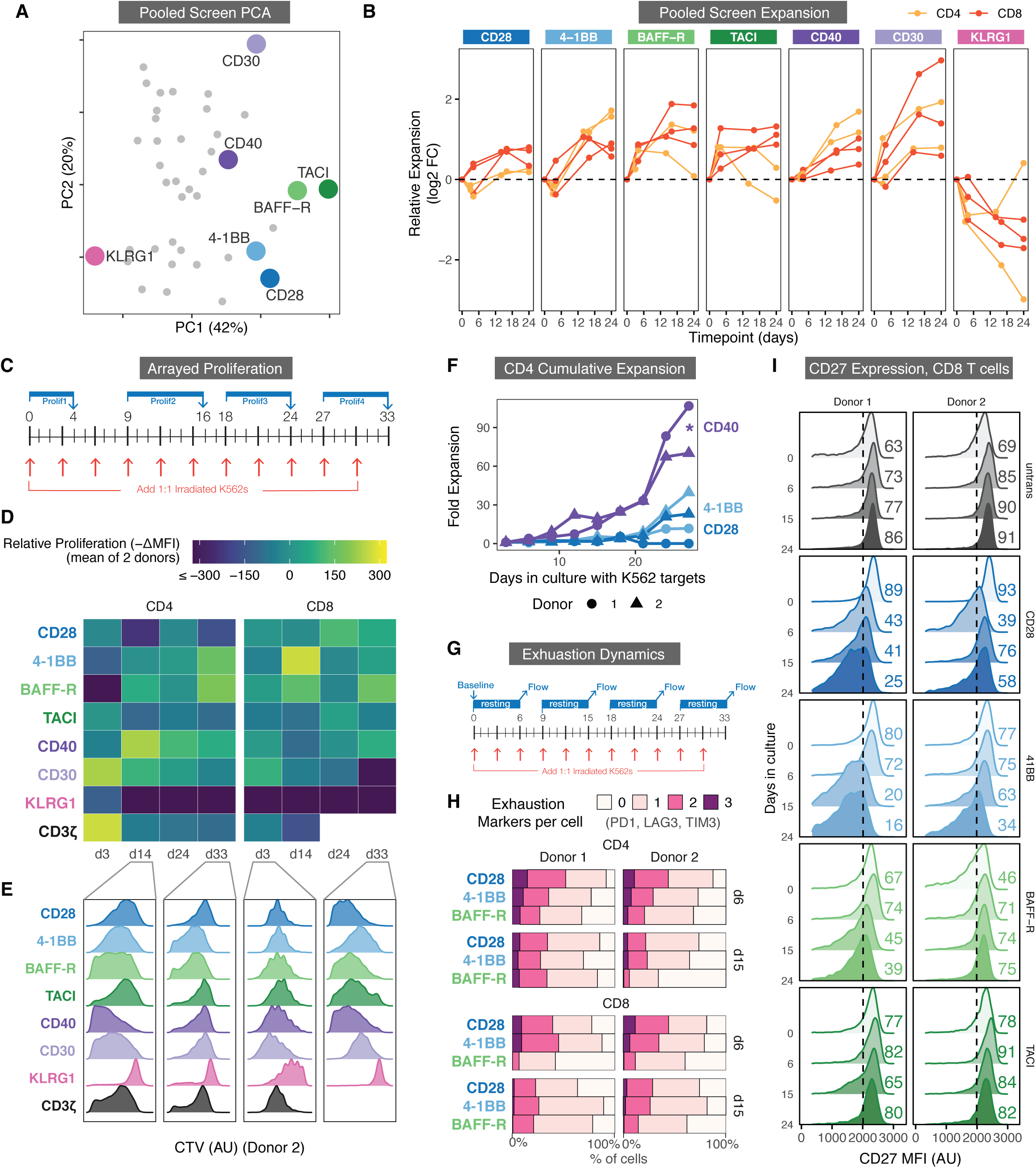
Novel set of cosignaling domains differentially affect proliferation, long-term expansion, markers of memory, and exhaustion. **A)** Principal components analysis (PCA) of pooled library screen cytokine, proliferation, expansion, and activation data for CD4s and CD8s, across both CD19+ and CD19- stimulation conditions, all donors, and all analysis timepoints. Chosen CARs are larger and shown in representative colors. **B)** Relative expansion of library members CD28, 4-1BB, BAFF-R, TACI, CD40, CD30, and KLRG1 over 24 days of repeated stimulation with irradiated CD19+ K562 cells within the pooled screen. Expansion was quantified by calculating the fold-change of the proportion of each CAR within the library at each timepoint (X axis) as compared to baseline relative to the average CAR within the pooled library. Measured in CD4 and CD8 primary human T cells individually in 2-3 biological replicates. **C)** Experimental timeline for arrayed proliferation assays (described in D and E). Primary human CD4 and CD8 T cells were separately transduced with CD28, 4-1BB, BAFF-R, TACI, CD40, CD30, KLRG1, or CD3ζ-only CARs. Purified CAR T cells were then stimulated 1:1 with irradiated CD19+ or CD19- K562 cells every three days. Proliferation was assessed via CTV every 9 days. **D)** Relative proliferation of each CAR quantified by the relative decrease in MFI (i.e., dilution of CTV dye) between two donors of CD4 or CD8 T cells. More proliferative cells are in yellow, less proliferative in dark blue. The X axis indicates the day the cells were stained. White boxes are representative of CAR T cells that stopped proliferating and dropped out of culture by that time point. **E)** Histograms of CTV staining of CD4 or CD8 CAR T cells in a representative donor. Data summarized in panel D. **F)** Quantification of the cumulative expansion of CD4 T cells engineered with either CD40 (purple), 4-1BB (light blue), or CD28 (dark blue) CARs and stimulated as described in panel C. Co-cultures were measured every three days starting on day 3 via flow cytometry and counting beads. Y axis measures cumulative fold-expansion every three days (see Methods). **G)** Cells were transduced and stimulated as described in panel C. Every 9 days in culture, cells were rested for 6 days without additional stimulation and assessed for surface expression of PD1, LAG3, and TIM3 (known markers for exhausted T cells). **H)** Number of CD8 CAR T cells expressing 0-3 of the exhaustion markers PD1, TIM3, LAG3 after day 6 and day 15 as described in G. All CARs, markers, and time points are shown in Figure S3A and S3B. **I)** CD27 surface expression on CD8 CAR T cells measured over 33 days in culture, with the same protocol as described in panel G. Percentage of CD27-high cells is shown for each CAR and day on the right, showing that CD28 and 4-1BB CARs lose CD27 expression over time compared to BAFF-R and TACI CARs. All CARs and time points are shown in Figure S3D.

There was no discernable clustering based on cell-type specific expression or protein family structure, aside from the aforementioned enrichment of TNF family members in the potent costimulatory group (Figure S2B). Notably, some domains which are known to be inhibitory in their natural contexts, such as PD1 and CTLA4, did not significantly reduce CAR activation or proliferation. While PD1 and CTLA4 have previously shown to be functional in an inhibitory CAR format *in trans (Fedorov and Themeli, 2013)*, all of the CARs in this work use the CD8 hinge and transmembrane domain and contain a C-terminal CD3ζ domain, which may significantly alter their signaling. However, some domains, such as KLRG1 and NKR-P1A, were able to significantly reduce proliferation and activation.

### A novel set of cosignaling domains differentially affect proliferation, long-term expansion, markers of memory, and exhaustion

As not all measurements can be easily performed in a pooled format, a subset of activating and inhibitory CARs were tested in greater depth. We included domains from each of the three subsets of potent costimulatory domains described above and selected those with enhanced resistance to exhaustion. These domains included BAFF-R (light green), TACI (dark green), CD40 (light purple), and CD30 (dark purple). Additionally, KLRG1 (pink) was included as the top potential inhibitory domain, as T cells expressing this CAR consistently demonstrated the lowest initial proliferation, activation, and cytokine production (Figure 2F). KLRG1 was also of interest because strong inhibitory cosignaling could be used to halt, dampen, or dynamically modulate T cell functions (Fedorov and Themeli, 2013). T cells expressing the various CARs demonstrated variable relative expansion dynamics within the pooled library, with those expressing BAFF-R, TACI, CD40, and CD30 exhibiting enhanced persistence by day 24 (Figure 3B). In addition, T cells expressing the chosen CARs also displayed a range of proliferative phenotypes in the absence of antigen, with CD30 showing the most antigen-independent expansion and CD40 showing pronounced contraction without antigen stimulation (Figure S2D). Finally, 4-1BB, CD28, and a CD3ζ first-generation CAR were included as benchmarks due to their clinical relevance and abundance of characterization within the literature (Kawalekar et al., 2016; Sun et al., 2020). All CARs were similar in their surface expression and T2A-GFP reporter fluorescence (Figure S2F).

An extended *in vitro* repetitive stimulation assay was performed individually on these 8 CARs to confirm their proliferative performance in CD4s and CD8 T cells. Proliferation was measured weekly using CTV over 33 days of repetitive antigen stimulation in two separate T cell donors (Figure 3C, S2E). Representative CTV dilution measurements for a single donor are shown alongside quantifications of the average change in MFI of the Cell Trace Violet stain (Figure 3D, 3E). These arrayed stimulations resulted in relative proliferations which were similar to the pooled screen, with CD28 demonstrating less proliferation in CD4s at later time points, CD40 generating strong proliferation in CD4s, 4-1BB and BAFF-R demonstrating stronger late-stage proliferation through day 33, and KLRG1 displaying dramatically less cell division overall (Figure 3D). Lastly, contrary to our pooled data, CD30 drove an initial burst of proliferation, primarily in CD4s, but this was not sustained in later weeks (Figure 3D, 3E).

In addition to measuring proliferation via CTV dilution, we counted the number of T cells in culture every 3 days prior to restimulation with additional K562 tumor cells via flow cytometry and cell-counting beads. We used these counts to calculate the overall cumulative expansion of each CAR. Similar to the relative expansion measurements in our pooled screen, this takes into account both proliferation and resistance to cell death. In CD4 T cells from both donors, 4-1BB had a higher degree of cumulative expansion and persistence than CD28, in line with the clinical findings that 4-1BB CAR Ts are better long-term proliferators *in vivo* and are more resistant to exhaustion than CD28 CAR Ts. However, we found over 33 days, CD4 CAR T cells with CD40 costimulation doubled at an average of 1.8x the rate of those with 4-1BB or CD28 (*p*=3.3×10^-3^), indicating, as in our pooled experiment, a heightened propensity for proliferation and resistance to cell death during prolonged antigen stimulation (Figure 3F).

A recent study strongly associated differences in mitochondrial metabolism with differences in long-term proliferation (Kawalekar et al., 2016). We used SCENITH, a method which uses oligomycin to inhibit mitochondrial oxidative phosphorylation (OXPHOS) (Argüello et al., 2020) to determine the relative contribution of OXPHOS to the overall metabolic output of the CAR T variants after 21 days in culture. As expected, the CD3-ζ-only CAR T demonstrated a distinctly low level of mitochondrial metabolism in both CD4s and CD8s, indicating increased dependence on glycolysis. 4-1BB and CD28 CAR Ts were biased towards OXPHOS or glycolytic metabolism respectively, as previously noted in the literature (Kawalekar et al., 2016). BAFF-R CAR T cells exhibited even higher mitochondrial dependence than 4-1BB after 21 days, in line with its long-term persistence in culture after repeated stimulations (Figure S2G).

### BAFF-R and TACI CARs retain markers linked to persistence and demonstrate delayed exhaustion marker expression

To determine the relationship between cell state and expansion of these CAR T variants, we measured the expression of several exhaustion markers (PD1, LAG3, TIM3, CD39) and differentiation markers (CD62L, CD45RO, CD45RA, CD27, CCR7) throughout 33 days of prolonged stimulation (Figures 3G, S3A-D). While the CARs showed relatively similar differentiation over time (Figure 3C), BAFF-R CAR Ts had a slower increase in exhaustion marker expression than 4-1BB and CD28, as shown by its overall lower number of exhaustion markers on days 6 and 15 in both CD4s and CD8s (Figure 3H, S3B). Additionally, both BAFF-R and TACI CD8 CAR T cells showed sustained CD27 expression over time, in contrast to CD28 and 4-1BB, in which CD27 expression progressively decreased (Figure 3I). We did not observe this trend in CD4s (Figure S3D). CD27 expression has been linked to CD8 T cell survival after extensive proliferation, and resistance to terminal effector differentiation and contraction (Carr et al., 2006; Dolfi et al., 2008; Hendriks et al., 2000, 2003). Finally, as seen in our proliferation assays, KLRG1 CAR T exhaustion and differentiation marker expression was most similar to untransduced T cells, suggesting that KLRG1 inhibits activation mediated by the CD3ζ domain, keeping a larger fraction of the population in a naive-like or memory state (Figure S3A, S3B, S3C, S3D).

### Comparison of cytokine secretion and *in vitro* toxicity across cosignaling domains

In addition to proliferation and persistence, we sought to measure differences in CAR T cytotoxic activity by both cytokine secretion and tumor cell killing ability. We measured cytokine production in CD4 T cells in two donors via intracellular staining after 1, 2, 3, 6 and 9 repeated stimulations in culture (Figure 4A, S4A). Most CARs exhibited maximal cytokine production on day 4, after the second antigen stimulation. While comparisons can be made at early time points, none of the CARs, including those containing the CD28 and 4-1BB costimulatory domains, produced significant amounts of IFNγ, IL2, or TNFα , as measured by intracellular staining after 3 or more *in vitro* stimulations.

**Figure 4:**
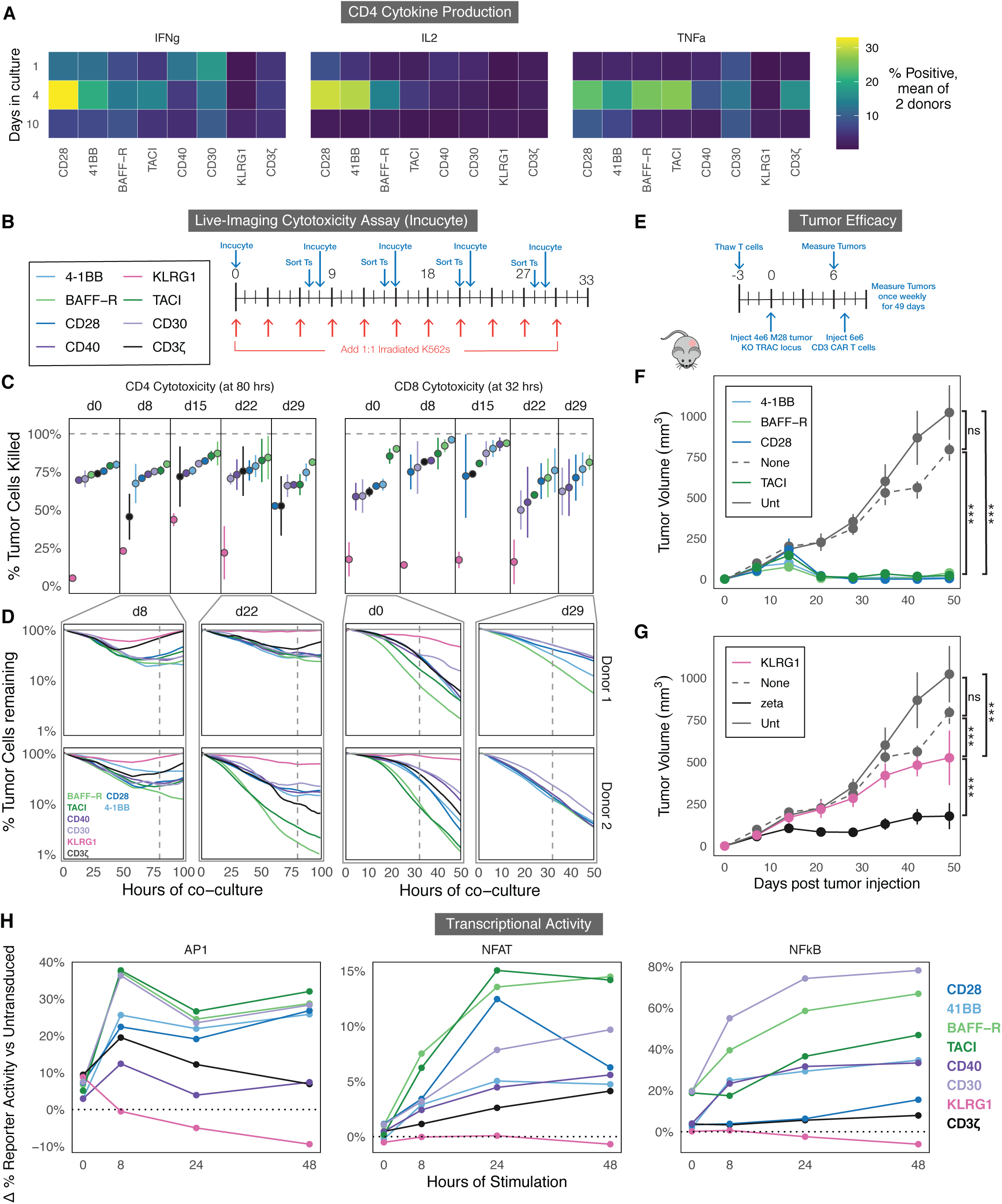
Comparison of cosignaling domains across measures of cytokine secretion, T cell signaling reporters, *in vitro* cytotoxicity, and *in vivo* solid tumor clearance. **A)** Mean cytokine production at 1, 4, and 10 days, measured across two donors by intracellular cytokine staining. Each CAR was stimulated with K562s as in Figure 3C and cytokines were accumulated intracellularly with brefeldin and monensin before staining and FACS (see Methods). The percentage of cytokine-positive cells of each type was averaged between the two donors. **B)** Experimental timeline for *in vitro* cytotoxicity assays. CD4 or CD8 primary human T cells were transduced with a CAR construct, selected by FACS, and exposed to repeated stimulations with irradiated K562s. Once weekly, a portion of T cells were purified from the co-culture via FACS and rested overnight. These T cells were then cultured 1:1 with mKate+ CD19+ K562s and imaged every 60 minutes via Incucyte for the next 3-5 days (see panels C and D). **C)** Cytotoxicity of CD4 (left) or CD8 (right) CAR T cells were quantified at 80 and 32 hours, respectively, by calculating the percentage of mKate+ tumor cells at each time point relative to a control condition with no T cells. CARs are ranked at each timepoint from least to most cytotoxic (left to right). **D)** Representative plots of Incucyte cytotoxicity assays in two donors at day 8 and day 22 for CD4s (left) and day 0 and day 29 for CD8s (right) plotting the percentage of mKate+ tumor cells remaining relative to a well with tumor cells but no T cells (grey). Vertical dashed lines indicate the time points analyzed in panel C. **E)** Experimental timeline for *in vivo* tumor models. We injected 4e6 M28 mesothelioma tumor cells subcutaneously into the flanks of NSG mice and, seven days later, transferred 6e6 engineered TRAC-knockout CAR T cells intravenously into the tail vein. Tumors were measured via caliper every 7 days for a total of 49 days. **F)** Tumors were either left untreated, treated with untransduced T cells, or treated with engineered CAR T cells. Tumor size was monitored over 49 days post tumor injection. CAR T cells containing the potent costimulatory domains are shown, compared to untransduced T cells and no T cell controls. **G)** Tumor size as in F, showing a CD3ζ-only control and a KLRG1 inhibitory CAR, compared to untransduced T cells and no T cells. **H)** Transcriptional activity reporter Jurkat cell lines for AP1, NFAT, and NFκB were transduced with each CAR and sorted within one log of GFP expression. The cells were stimulated with CD19+ K562s for 0, 8, 24, or 48 hours and then assessed for activity via flow cytometry. Percent transcription factor activity relative to untransduced reporter Jurkat cells is plotted on the Y axis. Resting and CD19- K562-stimulated CAR T cells are plotted in Figure S4E.

KLRG1 showed reduced IFNγ, IL2, and TNFα production as compared to CD3ζ-only CD4 CAR T cells at every time point. On the other end of the spectrum, CD28 CAR T cells produced a significantly higher level of IL2 and IFNγ during the first two weeks of stimulation, matching data from our pooled screens and in line with prior observations in the literature and in CAR T patients (Ying et al., 2019). High cytokine production within CAR T cells has been associated with higher rates of CRS during initial response and increased propensity for late stage T cell exhaustion. CD4 CAR T cells with BAFF-R and TACI costimulatory domains show a similar level of cytokine secretion to those with 4-1BB.

We next sought to longitudinally measure each CAR’s cytotoxic capabilities in culture after intervals of repetitive antigen stimulation up to one month. To directly measure cytotoxicity *in vitro*, we used the Incucyte live-cell imaging system, which allows for long-term imaging of fluorescently labelled tumor and T cell co-cultures inside of an incubator. At multiple timepoints after the repeated antigen stimulations, we FACS-sorted either CD4 or CD8 T cells from the co-culture with irradiated K562 tumor cells and let them rest overnight. The next day, we combined the sorted T cells with red-fluorescent, live K562 tumor cells at a 1:1 ratio and performed time-lapse live-cell microscopy on the co-culture to observe cell killing (Figure 4B). We then quantified the percentage of tumor cells killed after either 80 hours (CD4s) or 32 hours (CD8s) after different numbers of prior repetitive stimulations (Figure 4C). Killing dynamics from several representative live-imaging experiments for two donors are shown in Figure 4D as the percentage of K562 tumor cells killed over time.

While we observed differences in the cytotoxic capacity between the two human donors, BAFF-R and TACI consistently showed superior cytotoxicity relative to other CARs, both by rate of killing and by total percentage of tumor cells killed, especially after multiple rounds of prolonged antigen stimulation (Figure 4C, 4D). This was especially pronounced in CD4 T cells, where BAFF-R and TACI often ranked as the top two cytotoxic domains (Figure 4C). We repeated these assays with two additional donors in CD4s, comparing the top four domains (CD28, 4-1BB, BAFF-R, TACI), and confirmed that the latter two non-standard costimulatory domains showed equivalent or enhanced cell killing (Figure S4C).

Additionally, we saw that KLRG1 had drastically reduced cytotoxicity, often killing between 0-20% of K562 tumor cells, compared to the CD3ζ-only CAR, with which approximately 75% of tumor cells were killed at each timepoint (Figure 4C). Combined with the proliferation, exhaustion, and differentiation data, this supports the hypothesis that KLRG1 inhibits the CD3ζ domain and significantly dampens CAR T cell function.

### BAFF-R and TACI CARs clear an *in vivo* solid tumor model

To validate our observations *in vivo*, an established mesothelioma solid tumor model (M28) was used, which is known to produce durable tumors that require CAR persistence rather than rapid initial proliferation (Hyrenius-Wittsten et al., 2021). The slower growth dynamics of the M28 model give a longer window of time to determine clinical efficacy against solid tumors and the propensity for relapse. To utilize the M28 system, we exogenously expressed CD19 on M28 cells and sorted for cells with CD19 expression similar to the K562s. We injected NOD-*scid* IL2Rgamma^null^ (NSG) mice with 4×10^6^ M28 cells and 7 days later treated them with 6×10^6^ anti-CD19 CAR T cells. To compare the rejection dynamics and ensure CARs targeting the exogenously expressed CD19 antigen are comparable to CARs targeting the endogenously expressed ALPPL2 antigen, we set up a side-by-side *in vivo* experiment utilizing either anti-ALPPL2 4-1BB or anti-CD19 4-1BB CAR treatments. We saw no differences in tumor growth or rejection dynamics between CAR treatments targeting the two antigens (Figure S4D). Having confirmed similar tumor rejection between CARs targeting the engineered and natural ligands, we compared the different CAR costimulatory domains head-to-head in the M28 CD19 model (Figure 4E). We were unable to distinguish a difference between 4-1BB and CD28 CARs at the time points shown here despite the large amount of evidence from human studies that 4-1BB has increased long-term killing and persistence. Both 4-1BB and CD28, as well as our novel TACI and BAFF-R CAR treatments, exhibited similar tumor clearance and remission over 50 days (Figure 4F). Additionally, KLRG1 mirrored the results from the *in vitro* cytotoxicity assay and showed markedly increased tumor burden compared to all CARs, including the CD3ζ-only CAR, while showing modest efficacy compared to untransduced T cells (Figure 4G, S4E).

### Transcriptional reporters indicate differences in early signaling dynamics among the costimulatory domains

While a majority of our analyses indicated distinctions in CAR T cell phenotypes after prolonged periods of stimulation *in vitro* and *in vivo*, we sought to determine if there were early differences in signaling upon the initial activation of each CAR that could help to understand the mechanisms behind these phenotypic differences. We transduced each of the 8 CARs into three reporter Jurkat T cell systems that individually measured the transcriptional activity of activator protein 1 (AP-1), nuclear factor of activated T cells (NFAT), and nuclear factor kappa B (NFκB) (Hyrenius-Wittsten et al., 2021). We then stimulated these purified CAR-positive Jurkat reporters with CD19+ or CD19- K562 tumor cells for 8, 24, or 48 hours and measured their activity via flow cytometry (Figure 4H, Figure S4E). We observed significant differences between each CAR’s induction of transcription factor (TF) activity. KLRG1 CAR T cells had reduced activity for all three transcription factor reporters as compared to CD3ζ-only CAR T and untransduced T cells. Additionally, across all three reporters, BAFF-R and TACI showed both accelerated dynamics and a higher total percentage of cells with TF activity. This increase in TF reporter activity was most pronounced for NFκB, which is closely associated with TNF receptor signaling. As expected, we also saw a higher level of basal NFκB signaling from the TNF receptor family CARs, and a reduced basal AP1 signal from CD40, which correlates with its lack of tonic signaling (Figure S4E).

### Single-cell RNAseq and CITEseq characterize functional differences between novel CAR costimulatory domains

The marked differences between the CARs in the transcriptional reporter assay suggested that a deep and unbiased look into early transcriptomic signatures could explain their long-term functional differences in cytotoxicity, proliferation and exhaustion which we observed throughout our repetitive stimulation co-culture. Previous studies have used single-cell RNA sequencing (scRNA-seq) to compare CARs containing CD28 and 4-1BB domains, identifying differences in signaling, metabolism, and differentiation (Boroughs et al., 2020). To achieve a comprehensive understanding of the phenotypic landscape of CAR T cells incorporating these new signaling domains, we measured single cell RNA expression and a CITE-seq antibody panel of 75 proteins (Stoeckius et al., 2017) in order to map the unbiased transcriptome measurements onto well-studied T cell surface markers. We separately transduced each CAR into bulk CD3 T cells from two PBMC donors and performed 10x Chromium 3’ v3 scRNAseq after four days in culture, either with or without stimulation provided by irradiated CD19+ K562s. (Figure 5A).

**Figure 5:**
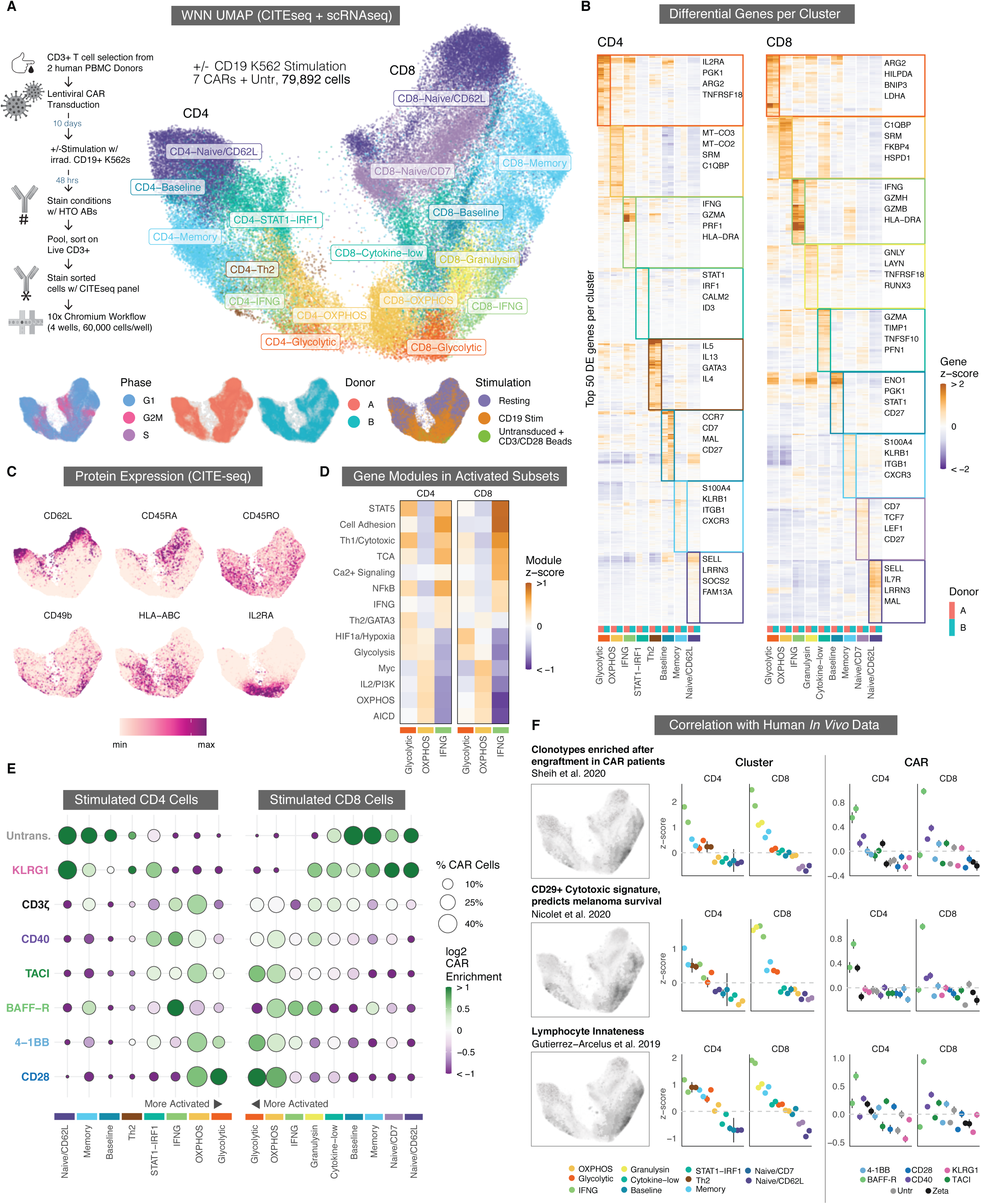
Single-cell RNAseq and CITEseq characterize functional differences between novel CAR costimulatory domains. **A)** Weighted-nearest neighbor (WNN) Uniform Manifold Approximation and Projection (UMAP) embedding of scRNAseq and CITEseq data from stimulated and resting CAR T cells from two donors, as described in the experimental timeline on the left. UMAP separates into well-defined CD4 and CD8 lobes (left and right sides of the embedding). Main UMAP is separated by color into 8 CD4 and 9 CD8 phenotypic clusters. Bottom inset: UMAPs are colored by cell cycle phase, donor identity, and stimulated vs resting cells. Figure S5A shows a version of this UMAP faceted separately for each CAR and stimulation condition. **B)** Expression heatmap of differentially expressed genes in all CD4 and CD8 clusters. For each cluster, the top 50 differentially expressed genes are shown, and are ordered by hierarchical clustering of the pseudo-bulk expression Z-scores for all clusters and both donors. Genes in the top 50 for multiple clusters are only included for the cluster with the highest Z-score. For each cluster, four genes within the top 50 that are representative of the overall phenotype of the cluster are plotted in the right-hand boxes. **C)** UMAP plots show relative CITEseq expression for the surface expression of 6 markers associated with T cell differentiation and activation. **D)** Mean z-scores for MSigDB gene modules associated with various aspects of T cell activation, metabolism, and signaling among the three major activated phenotypic clusters in CD4 and CD8 T cells. **E)** Enrichment of stimulated CAR T cells containing different cosignaling domains within each phenotypic cluster. The size of each dot corresponds to the percentage of stimulated CAR - T cells in a cluster and with a costimulatory domain. The color of each dot corresponds to the log-2 fold cluster enrichment or depletion of that CAR relative to others. Clusters are arranged with the most activated at the center to correspond to the UMAP in panel A. Similar plots for resting cells and a per-donor breakdown are in Figures S5C and S5E, respectively. **F)** Correlation of T cell gene signatures indicative of enhanced CAR T engraftment (top), melanoma survival (middle), and lymphocyte innateness transcriptional signature (bottom), with phenotypic clusters in CD4 and CD8 CAR T cells (middle column) or with CARs containing different costimulatory domains (right column). Cluster and CAR colors match those in panel E. The two dots per group correspond to donors A and B.

We used a weighted-nearest neighbor graph-based clustering approach to combine both the CITE-seq and scRNAseq data across 79,892 cells, followed by UMAP dimensionality reduction (Hao et al., 2021). This combined protein and RNA embedding separated the cells into well-defined CD4 and CD8 lobes (left and right), with resting cells at the outer edges and stimulated CD4 and CD8 cells in the center bottom, in distinct but adjacent regions (Figures 5A, S5A), suggesting that CD4 and CD8 CAR T cells converge towards a more similar activated phenotype after CAR stimulation. Additionally, while the cells were grouped into 8 CD8 clusters and 9 CD4 clusters (Figure 5A, 5B), we noticed a pronounced mirroring of transcriptional programs across 5 pairs of clusters between CD4s and CD8s (Naive/CD62L, Memory, IFNG, OXPHOS, and Glycolytic), and thus describe these clusters with matching labels (Figure S5B). Resting CAR T cells were found almost entirely in naive-like (Naive/CD62L and Naive/CD7) and memory clusters, while stimulated cells fell into three main clusters with differing cytotoxic and metabolic transcriptional and surface protein signatures (discussed below), as well as a few other clusters with phenotypes in between naive-like and fully activated (Figure 5A-C).

### Activated CAR T cells fall into 3 distinct clusters shared among CD4s and CD8s

In both CD4 and CD8 T cells, activated CAR T domains segregate into 3 distinct clusters we named Glycolytic, OXPHOS, and IFNG, which differ in the expression of several important metabolic genes and signaling pathways (Figure 5B-5D). Cells in the Glycolytic cluster differentially upregulate genes involved in aerobic glycolysis (*ARG2*, *PGK1*, *LDHA*), the HIF1a pathway (*SLC2A3*, *ENO1*, *ALDOC*), mitochondrial autophagy (*BNIP3*), protein markers involved in costimulation and T cell activation (*IL2RA*, *CD69*), and inhibitory receptors (*PD1*, *CTLA4*) (Figure 5B, 5D, S5E). Cells in the OXPHOS cluster upregulate genes involved in oxidative phosphorylation and arginine metabolism (*SRM*, *C1QBP*, *ATP5MC3*, *MT-CO3*), as well as the Myc, MTOR, and PI3K pathways (Figure 5D). The third activated cluster, which we named IFNG, has a strong cytotoxic expression signature, expressing a high level of IFNG transcript, as well as multiple granzymes and cytotoxic molecules including *GZMB*, *GZMK*, *GZMH*, *NKG7*, and PRF1. Both CD4 and CD8 IFNG cells also abundantly express both RNA and protein for MHC class II and CD74, an MHC class II chaperone, as well as a variety of integrins and chemokine receptors including *ITGB2* (CD29), *ITGA2* (CD49b), *ITGA4* (CD49d), *CXCR3*, and *CCR5*. The IFNG cluster is more similar to Glycolytic than OXPHOS but shows differential and overall lower expression of various inhibitory and activation protein markers, including IL2RA, OX40, 4-1BB, PD1, and GITR, while expressing a higher level of memory markers, including CD95 and CD45RO (Figure 5C).

Having identified a variety of activation states and transcriptional signatures across CAR clusters, we next sought to identify those which were differentially enriched among costimulatory domains (Figure 5E, Figure S5A, S5C, S5E). While all 5 costimulatory domains are present in the three most activated clusters, CARs containing the CD28 costimulatory domain were enriched in the CD4 and CD8 Glycolytic clusters, while the BAFF-R CAR is particularly enriched in the IFNG cytotoxic clusters. BAFF-R is also the most enriched in the Memory cluster after stimulation, indicating that a larger proportion of CAR T cells containing this domain remained in a less activated and less differentiated state after CAR stimulation. This divergence in BAFF-R and CD28 enrichment was observed in both donors (Figure S5E).

### IFNG cluster matches multiple signatures of improved clinical response, CAR engraftment

Neither mouse xenografts nor *in vitro* assays are the ideal metric with which to compare efficacy of adoptive transfer therapy within patients, but the unbiased and high-dimensional data acquired from single cell sequencing offers an opportunity to identify transcriptional signatures which correlate with positive clinical outcomes in patients. We found that the cytotoxic IFNG cluster closely matches gene signatures associated with TIL and CAR efficacy identified in two recent studies. A study of clonal kinetics in patients undergoing CAR T immunotherapy identified a gene signature enriched in CAR T clones which preferentially expanded (IRF, increased relative frequency) in patients 1-2 weeks after infusion (Sheih et al., 2020). We found that this IRF gene signature showed a distinct and significant overlap with our CD4 and CD8 IFNG clusters (p = 1×10^-9^), and that the BAFF-R CAR showed particular enrichment for this CAR expansion signature (Figure 5F). A second study by Nicolet and colleagues identified a T cell expression signature which was shown to result in enhanced IFNγ secretion and increased survival in melanoma patients (Nicolet et al., 2020). Their study associated this cytotoxic gene signature with the integrin CD29. In the IFNG cluster, and in BAFF-R CAR T cells in particular, we see a marked upregulation of CD29 as well as CD49d, which together form the heterodimeric VLA-2 integrin complex, which is associated with more potent effector memory T cell responses (Figure S5D, S5F) (Kassiotis et al., 2006; Yan et al., 2008).

In addition to the increased expression of MHC genes and integrins, the IFNG and Memory clusters also express several receptors more typical of NK cells, including *KLRB1*, which encodes CD161. CD161 is expressed by a subset of T cells with characteristic tissue homing (Billerbeck et al., 2010) and increased cytotoxicity (Fergusson et al., 2014), and recent work has shown that CD8+CD161+ T cells define a potent effector memory subset with enhanced CAR T cell efficacy (Konduri et al., 2021). To explore this further, we turned to a recent study which identified a continuous gene expression gradient of lymphocyte innateness from T cells to NK cells (Gutierrez-Arcelus et al., 2018). This innate lymphocyte gene expression program largely overlaps with the transcripts enriched in our IFNG and Memory clusters, transcripts enriched in the activated BAFF-R CAR, and the clinical CAR T engraftment and CD29/IFNγ expression signatures we identified from recent literature (Figure 5F, S5F). The unexpected overlap among these disparate data suggests a linkage between innate-like gene expression and beneficial cytotoxic CAR phenotypes. Overall, we show that these novel CAR signaling domains promote altered cell states, including some that are associated with potentially beneficial anti-cancer responses.

## DISCUSSION

Costimulation is essential for long-term T cell proliferation, differentiation, survival, and it is known to strongly enhance CAR T cell efficacy. While canonical T cell costimulatory domains like 4-1BB and CD28 are well-studied, a wide variety of known cosignaling domains have yet to be tested in CARs, and the extent to which signaling domains from other cell types can act upon T cells is poorly understood. Here, we identified several novel cosignaling domains through pooled library screens in primary human T cells that enhance persistence or cytotoxicity over those used in the current generation of FDA-approved CARs. By screening 40 domains, we show the breadth of the cosignaling landscape within primary human CAR T cells. We observed that many of the most potent costimulatory domains measured belonged to the TNF receptor family, to which 4-1BB also belongs. 4-1BB signaling is known to increase T cell persistence and late stage proliferation relative to CD28 (Weinkove et al., 2019), and our screen shows that this property extends to several other members of the TNF receptor family, especially CD40 and BAFF-R. Based on the results of our pooled screens, we selected 4 new stimulatory CARs from the TNF receptor family, as well as a novel inhibitory CAR, and performed comprehensive characterization of their proliferation, persistence, differentiation, exhaustion, cytokine production, and cytotoxicity both *in vitro* and *in vivo* over a 4-5 week period. Characterizing these domains separately in CD4s and CD8s allowed us to identify effects specific to each cell type. We found that the BAFF-R and TACI signaling domains had heightened cytotoxicity, CD40 had heightened persistence in CD4s, and KLRG1 can potently inhibit ITAM-based signaling, preventing many hallmark features of T cell activation and differentiation.

To identify differences in gene expression elicited by our curated set of costimulatory domains, we explored each CAR’s transcriptomic profile and surface protein expression via scRNAseq and CITEseq both before and after antigen exposure. We observed distinct differences in gene and protein expression between the 8 CARs and partitioned cells into clusters based on their degree of differentiation, activation, cytotoxicity, and metabolism. We observed a distinctive mirroring of many of these clusters across the CD4-CD8 axis. Activated CAR T cells fell into three distinct clusters which differed in their expression of key metabolic and cytotoxic genes. We named these clusters Glycolytic, OXPHOS, and IFNG. The IFNG cluster was significantly enriched for CARs containing the BAFF-R costimulatory domain, which also showed strong cytotoxic and proliferative performance in our prior *in vitro* and *in vivo* assays. Next, we found significant overlap between this IFNG cluster and gene signatures from several other recent studies of 4-1BB CAR infusion products and TIL efficacy in cancer patients, suggesting that this signature correlates with higher engraftment, persistence, and tumor rejection. Compared to 4-1BB, BAFF-R CAR-T cells are approximately two-fold enriched (3x in CD4s, 1.5x in CD8s) for this gene signature, suggesting that a BAFF-R CAR infusion product would have a much larger fraction of cells in this hyper-effective state, potentially resulting in improved patient outcomes. Most surprisingly, this gene signature, as well as those of the other correlated clinical studies, maps onto an innate lymphocyte program more akin to iNKTs, γδ T cells, and NK cells than to αβ T cells. While this association remains correlative, and the relationship between CAR efficacy and NK-like gene expression is complex(Fergusson et al., 2014; Konduri et al., 2021; Mathewson et al., 2021), it suggests an exciting intersection between different CAR costimulatory signaling phenotypes and the promising efficacy of CAR-NK cells(Xie et al., 2020).

Although ‘CAR Pooling’ allows us to directly compare an unprecedented number of signaling domains, discrepancies can arise between the performance of domains in a pooled versus an arrayed setting. For instance, we observed robust long-term expansion of CD30 in the pooled screens but more rapid exhaustion in the follow-up arrayed screen. This could be potentially due to paracrine signals or cytokine production by neighboring CAR T cells in the pool. Subsequent arrayed testing of CARs can thus be critical to rule out paracrine effects. Another strategy may be to buffer or normalize paracrine effects by combining the library with a larger fraction of untransduced or first-generation CD3ζ T cells. Additionally, we also observed some differences between *in vitro* and *in vivo* performance, such as the lack of *in vivo* anti-tumor efficacy for CD40. To address such discrepancies, pooled screening of CAR T cell expansion and persistence could also be performed directly *in vivo.* Measurements of cytotoxicity (via cytokine production, granzymes, or CD107a) could also be performed *ex vivo* after intratumoral expansion of the library.

Although we are primarily concerned here with identifying improved costimulatory domains, it is important to note that the novel KLRG1 CAR is not simply a negative control but a potential tool to control CAR activity. Such a tool could be useful for reducing cytokine release syndrome, abrogating on-target off-tumor activation, and even generating post-translational oscillations in CAR activity that would mimic natural TCR signaling dynamics. Pooled screens can thus be employed not simply to find the best costimulatory domain but rather as a discovery tool to understand how different receptor signals alter T cell biology and how these signals can be modulated to manipulate T cell function beyond optimizing a single chimeric antigen receptor.

Much of the existing literature tends toward a one-dimensional comparison of CARs as more or less effective overall, while we instead observed that individual CARs often excelled within specific assays or when expressed in different T cell types. This multidimensionality of costimulation was somewhat unexpected, reinforcing that what is usually referred to as a monolithic ‘signal 2’ is instead a variety of heterogeneous pathways, sometimes in opposition, which can be individually tuned to optimize different aspects of T cell function. This also suggests that future engineering could isolate individual signaling motifs to enhance specific T cell phenotypes and create synthetic combinatorial domains which are optimized for specific ScFvs, tumor types, or to better combat the functional deficiencies seen in CARs for solid tumors. Finally, in addition to engineering better therapeutics, future high-throughput studies will help to elucidate the ‘design rules’ for synthetic receptors and signaling motifs, generate powerful new tools to manipulate T cells for basic immunology research, and lead us to a greater understanding of T cell differentiation, development, and immune-related disease.

## ACKNOWLEDGEMENTS

K.T.R. is funded by the Parker Institute for Cancer Immunotherapy, the UCSF Helen Diller Family Comprehensive Cancer Center, the Chan Zuckerberg Biohub, an NIH Director’s New Innovator Award (DP2 CA239143), Cancer Research UK, and the Kleberg Foundation. A.M is funded by Parker Institute for Cancer Immunotherapy, PICI, Burroughs Wellcome Fund, Career Award for Medical Scientists, The Cancer Research Institute, and a Lloyd J. Old STAR Grant. D.B.G. is supported by a Postdoctoral Fellowship from the Jane Coffin Childs Memorial Fund. C.S.A. is supported by a Diabetes, Endocrinology & Metabolism Training Grant, an NIH NIAD Training Grant, and a GWIS National Fellowship. C.J.Y. is supported by the NIH grants R01AR071522, R01AI136972, R01HG011239, and the Chan Zuckerberg Initiative, and is an investigator at the Chan Zuckerberg Biohub and is a member of the Parker Institute for Cancer Immunotherapy (PICI). We acknowledge the PFCC for assistance in generating Flow Cytometry data. Research reported here was supported in part by the DRC Center Grant NIH P30 DK063720.

## AUTHOR CONTRIBUTIONS

Names are listed alphabetically within each category. Conceptualization: C.S.A, J.A.B., D.B.G, A.M., K.T.R.; Methodology: C.S.A, D.B.G, K.K., K.T.R.; Software: D.B.G., K.G., K.K., N.P.; Investigation - Pooled Screens: C.S.A, D.B.G, K.K., E.P.; Investigation - Arrayed Screens: C.S.A, D.B.G., K.K.; Investigation - In Vivo: C.S.A, D.B.G., J.G., K.K.; Investigation - scRNAseq: C.S.A., D.B.G, B.H., D.L.; Writing Original Draft: C.S.A, D.B.G; Writing - Review & Editing: C.S.A, J.A.B., D.B.G, A.M., K.T.R.; Visualization: C.S.A, D.B.G, K.G., K.K.; Funding Acquisition: J.A.B., A.M., K.T.R., J.Y.; Supervision: J.A.B., A.M., K.T.R.

## METHODS

### Library Construction

Costimulatory intracellular domains (ICDs) were synthesized as gBlock gene fragments from Integrated DNA Technologies. These fragments were amplified with common primers containing homology to an E. coli cloning backbone, and the PCR products were individually cloned into the backbone in 96-well plates and colonies were sequence-verified after cloning using Sanger sequencing. Each cloned costimulatory domain was cultured and miniprepped separately and then pooled at 1:1 molar ratio. A fragment containing an SFFV promoter and N-terminal portion of the CAR up until the costimulatory domain was inserted in front of the pooled ICD plasmid library using Golden Gate Assembly and electroporation. This completed CAR cloning plasmid was then digested to separate the promoter, CAR, and a downstream GFP marker from the backbone. The digested plasmid library was finally gel-extracted and inserted via restriction cloning and electroporation into a digested and gel-extracted pHRSIN lentiviral backbone. We optimized the cloning procedure to ensure proper coverage of the library, with at least 1000 CFUs per library member, on average, at each pooled electroporation step.

### CellTrace Violet (CTV) dye

T cells were taken from culture, resuspended, and washed with PBS. We resuspended the culture to 1e6 cells/mL of a 5µM solution of Cell Trace Violet (CTV) in PBS and ncubated at room temperature for twenty minutes in the dark. We then added 5mL of T cell media on top for every 1e6 cells that were stained and incubated another 10 minutes in the dark. Then, cells were pelleted via centrifugation at 500g for 5 minutes and resuspended and plated in T cell media at 1e6 cells/mL. To stimulate proliferation, we plated an equal amount of K562s relative to the total number of T cells in each well, with or without CD19 expression, at a final concentration of 0.5e6 K562s/mL and 0.5e6 T cells/mL. We assessed the proliferation 3 days after stimulation for CD4s via flow cytometry. We restimulated T cells with an additional dose of K562s 3 days after the initial stimulation. To determine proliferation after prolonged co-culture, rather than restaining, we kept a proportion of the cells separate in culture and stained on day 9 (CTV2), day 18 (CTV3), or day 27 (CTV4) post-initial stimulation. We assayed proliferation on day 16 (CTV2), day 24 (CTV3), or day 33 (CTV4) via flow cytometry.

### CD69

To determine the degree of activation of each CAR we stimulated antigen-naive T cells with K562s, either with or without CD19 expression, in a 1:1 ratio. 24 hours after we plated the co-culture we centrifuged the cells at 500g for 5 minutes, washed twice into flow buffer (PBS + 2% FBS), and stained with anti CD69 antibodies at 4°C for 20 minutes. We washed the cells twice with flow buffer and ran on a flow cytometer.

### Cytokines

To determine the degree of cytokine production upon activation of each CAR, we stimulated antigen-naive T cells with K562s, either with or without CD19 expression, in a 1:1 ratio. 3 days after, we replated the co-culture with a secondary bolus of K562s (see the above method). We then added 2x Brefeldin A 12 hours later for an additional 6 hours. We centrifuged the cells at 500g for 5 minutes, washed twice into flow buffer (PBS + 2% FBS), stained with anti-CD4 or - CD8 antibodies at 4C for 20 minutes. We washed the cells twice with flow buffer and added 100uL of fixative (50uL of flow buffer + 50 uL of Invitrogen IC fix) to each well. We incubated at room temperature for 1 hour in the dark. After fixation we spun the cells at 600g for 5 minutes and resuspended in Cytolast for continued staining the next day. To permeabilize the cells we added 200uL of 1x Permeabilization buffer to each well, immediately spun at 600g for 5 minutes, and stained for intracellular antigens with anti-IL-2, -TNF□, and -IFNγ antibodies diluted in perm buffer. We stained in 50uL at room temperature for 30 minutes in the dark. We then washed twice with Permeabilization buffer and ran it on a flow cytometer.

### Incucyte

50 µL 5 µg/mL of fibronectin was dispensed to each utilized well of a 96-well plate. Plate was incubated for 60 minutes at room temperature, and fibronectin was removed, followed by another 60 minute incubation at room temperature. Both CAR T cells and live K562 target cells (either expressing mKate and CD19 or only mKate) were spun down and resuspend in Jurkat media + 30 U/mL IL-2; Jurkat media (RPMI-1640 medium + 10% FBS + 1% PenStrep + 1X Glutamax) has less fluorescence than media based on X-VIVO-15. Cells were counted and diluted to 0.25e6/mL each, and 100 µL of each (T cell and Targets) was added to each well for a final assay volume of 200 µL. Each condition was done in duplicate so long as sufficient cells were available. We allowed plates to settle at room temperature for 30 minutes before beginning the incucyte assay. Images were taken every 60 minutes using the Incucyte software over the course of the experiments (see relevant figures for total assay times, which varied between conditions).

### Splitting/Plating

Blood samples in the form of leukopaks were obtained from healthy male and female volunteers through STEMCELL Technologies. T cells were isolated via a CD4 or CD8 negative selection kit and frozen. We stimulated the T cells 24 hours after thawing with 25uL of CD3/CD28 beads (Thermo Fisher Dynabeads) per 1e6 T cells. Concentrated lentivirus was added 48 hours after thawing to reach a transduction rate of under 15% for the pooled library experiments and between 30-50% for the arrayed screens. Virus was removed within 18 hours of addition and cells were expanded. Five days after thawing, the beads were removed via magnetic separation and cells were sorted for GFP expression at least half a log higher than the negative population and spanning no more than a log in MFI. Cells were plated at 0.5e6 cells/mL and split every three days to this density until 10-14 days after thawing.

For the pooled experiments, we started stimulation on day 10 and for the arrayed experiments we started stimulation on day 14. To stimulate the T cells we combined them 1:1 with irradiated K562s (see next method) that either expressed or did not express surface human CD19 and plated at a density of 0.5e6 T cells/mL. Every three days we spun, resuspended, and counted the cultures to split the T cells and add more irradiated K562s. We counted the T cells by adding an aliquot of the resuspended culture to CountBright beads and calculating the number of T cells through analysis on the BD X-50 Flow Cytometer. The cultures were restimulated with the addition of additional irradiated K562s 1:1 to total T cells in each culture. They were replated at the density of 0.5e6 T cells/mL. This was repeated for a total of 3-33 days.

Media used was X-VIVO 15 + 5% hAb serum + 10mM NAC neutralized with 1N NaOH + 0.5% pen/strep + 1X beta-mercaptoethanol.

### Irradiation of Ks

Live K562 cells (ATCC® CCL-243™) were grown up in T182 flasks until confluent. Cells were resuspended to 10e6/mL on ice and irradiated using a Cesium-137 irradiator for 20 minutes (∼200 rad/min) in a 50mL falcon for a total dose of approximately 4,000 Rads. Cells were then aliquoted and frozen in IMDM media containing 10% DMSO and 10% FBS in liquid nitrogen until needed in the protocol.

### DNA extraction/sequencing

After fluorescence activated cell sorting assays, after in vitro growth with target cells, or after transduction as a library abundance baseline measure, T cells containing the CAR library were spun down into pellets, supernatant removed, and frozen at -80°C. Subsequently, genomic DNA was prepared from cells using either the Machery-Nagel Nucleospin Tissue XS column, Machery-Nagel Nucleospin column, or the Nucleospin 96 Tissue extraction plate. Manufacturer protocols were followed except for the addition of 10 µg polyadenylated RNA to each sample to increase yield.

After gDNA prep, Picogreen and a plate-based fluorescence reader were used to quantify the extracted genomic DNA. Initial PCR amplification of the costimulatory domain region from the different samples (PCR1) were done in 3 batches with differing numbers of cycles (12, 16 or 22 cycles) depending on the genomic DNA concentration. PCRs were performed with Takara ExTAQ to allow for maximum template concentration to be used in the PCR reaction. Reactions were done in 70 µL with between 200 and 1000 ng of DNA used as template depending on the batch as described above.

For the subsequent PCR to add Illumina barcodes and adapters to the products (PCR2), all products from PCR1 were diluted 15x and 25 µL of template was used in a 50 µL reaction with Takara ExTAQ. Different forward and reverse primers were used for each sample for PCR2 to add unique custom Illumina I5 and I7 barcode sequences to each sample.

Finally, PCR2 products were again quantified using Picogreen in a plate-based fluorescence reader. These products were pooled at 1:1 molar ratio, diluted, loaded, and run on a MiniSeq 2×150 cycle cartridge using the standard manufacturer protocols.

### Sequencing Analysis

After demultiplexing, CAR costimulatory domain sequences in FASTQ format for each sample were adapter-trimmed, sorted, deduplicated, and aligned using custom python scripts and BWA-mem. These alignments were then converted into count tables and analyzed using DESeq2 and custom R scripts (https://github.com/dbgoodman/tcsl-lenti).

### scRNAseq: T Cell Purification and Transduction

Experiments for each donor were performed separately on different days. CD3 T cells were purified from PBMCs extracted from either leukopacks or TRIMA residuals, as described in more detail above. CARs were then lentivirally transduced into bulk CD3s, using the same methods as previously described, and five days after transduction, T cells were FACS-sorted five days after transduction based on GFP marker expression based on a range of a one log away from the mean expression across all constructs. Untransduced T cells were not sorted.

### scRNAseq: T Cell Stimulation and 10x Prep

Five days after sorting (total of 10 days after transduction), 2e6 cells were plated with either 2e6 irradiated CD19+ K562 cells (described above) or replated in fresh media without K562 cells, at a density of 1e6 T cells/mL, for a total of 18 conditions per T cell donor: 6 second-generation CARs (excluding CD30), a CD3ζ only CAR, and untransduced T cells, both with and without K562 co-culture. For the second donor, a CD3/CD28 Dynabead-stimulation condition was also performed with untransduced cells, following instructions from the manufacturer. A K562-only control sample was also generated for both donors. After co-culture for 48 hours, cells in each condition were individually counted and stained CD3, DRAQ7, and with unique combinations of two TotalSeq-B hashtag antibodies (Biolegend) following the manufacturer’s instructions. After staining, the cells were pooled at approximately equal ratios based on counts taken prior to staining. The pooled cells were then sorted based on FSC/SSC, CD3, GFP, and DRAQ-7-negative gates to remove dead cells and irradiated K562s. Finally, the sorted cells were stained using the BD Bioscience ABseq antibody panel and loaded and processed via the standard Chromium V3 3’ sequencing pipeline. Two lanes were loaded per donor at 60,000 cells per lane. Cells were able to be loaded at this high density because doublets could be identified based on HTO barcode collisions between samples. One 200-cycle Novaseq S4 lane of Illumina sequencing was performed per donor (approximately 6e9 reads in total).

### scRNAseq: Data loading and cleaning

UMI counts were generated using CellRanger v5.0.1 using the standard settings and imported into Seurat v4, R v4.0.4, Rstudio v1.4. Barcode combinations were deconvolved using custom scripts and potential doublets were identified and removed based on barcode collisions (https://github.com/dbgoodman/tcsl-lenti). CD4 and CD8 cells were identified based on RNA and ADT expression of CD4/CD8, and double-positive cells were removed from consideration. Cells whose UMIs came from mitochondrial genes greater than 25% of the time were removed. A total of 79,892 cells remained after removing potential doublets and cells enriched for mitochondrial RNA.

### scRNAseq: Multidonor and multimodal data integration and clustering

Generation of UMAP embedding on both RNAseq and CITEseq data was performed using Seurat v4’s SCTransform and FindMultiModalNeighbors functions. Regression was done on cell cycle genes (Seurat’s *cc.genes*). Then, FindClusters was used to generate initial clusters with algorithm 3, cluster resolution 1.3. For initial data exploration purposes, this UMAP embedding and clustering procedure was performed separately for activated CD4s, activated CD8s, all CD4s, all CD8s, and finally with all cells (the latter generating the embedding shown in Figure 5A). Initial clusters were identified, then categorized and consolidated into the final clusters in Figure 5A, based on curation of the most differentially expressed genes and proteins, an extensive search of the literature, using the VISION tool, GSEA and mSigDB, and by hierarchical clustering of average gene expression across clusters.

---

**Figure S1:**
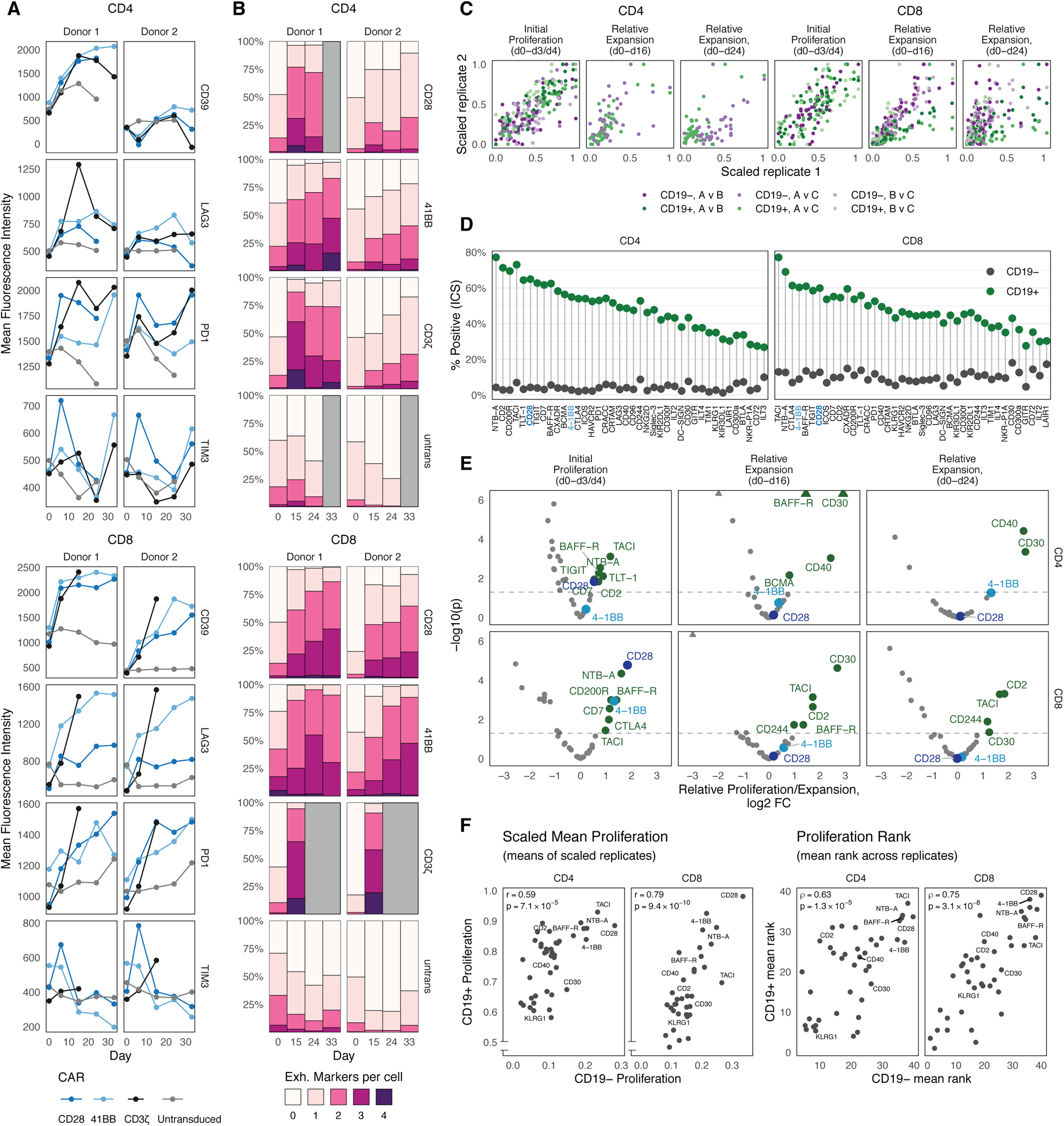
Repetitive stimulation reproducibly induces exhaustion and causes differential proliferation and expansion in a library of CAR-T variants. **A)** Mean fluorescence intensity (MFI) for four cell surface markers of exhaustion (CD39, LAG3, PD1, TIM3) in anti-CD19 CAR T cells generated from two donors measured after repeated stimulation with CD19-expressing irradiated K562s. CARs contained either 4-1BB or CD28 costimulation domains, no costimulatory domain (i.e., CD3ζ only), or were untransduced T cells from the same donor as a control. **B)** Aggregated measurements for S1A, displayed as the percentage of cells expressing 0-4 exhaustion markers, for each donor, timepoint, and CAR. **C)** Correlation between replicates across the pooled screens for initial proliferation on day 3 or 4 (d3/d4) and relative expansion within the pool after two and three weeks, shown separately for CD4 and CD8 T cells. Replicate A is Donor 1, Replicate B is Donor 2, and C is a second replicate of the library in Donor 2 that was run subsequently. Measurements within each replicate were individually rescaled from 0 to 1, with 0 being the least proliferative/expanded and 1 being the most proliferative/expanded. **D)** FlowSeq measurement of the percentage of CD69+ cells for each CAR library domain in both CD4 and CD8 cells, 18 hours after the addition of irradiated K562s either with or without CD19. Cells are ranked based on the difference in percentage of CD69+ cells between CD19+ and CD19- conditions. **E)** Volcano plots showing the relative proliferation or expansion (according to panel labels) of CD4 or CD8 T cells expressing CARs containing different costimulatory domains, during the repetitive stimulation assay with CD19+ K562s. The X axis shows the calculated difference in log2-fold proliferation/expansion, while the Y axis shows the associated adjusted P-value, as calculated by the DESeq2 algorithm. **F)** A comparison of CAR T cell proliferation from d0-d3 across the library with and without CD19 stimulation. The left plots show the scaled proliferation averaged over each replicate (as in C) but retain the differences in relative proliferation between CD19- and CD19+ conditions, which were measured simultaneously in our FlowSeq CTV assay. The right plots show the mean CAR rankings separately for the CD19- and CD19+ conditions. The top-performing potent costimulatory CARs from Figure 2E are labeled. On the left, the Y axis is truncated due to the higher relative proliferation in the CD19+ condition.

**Figure S2:**
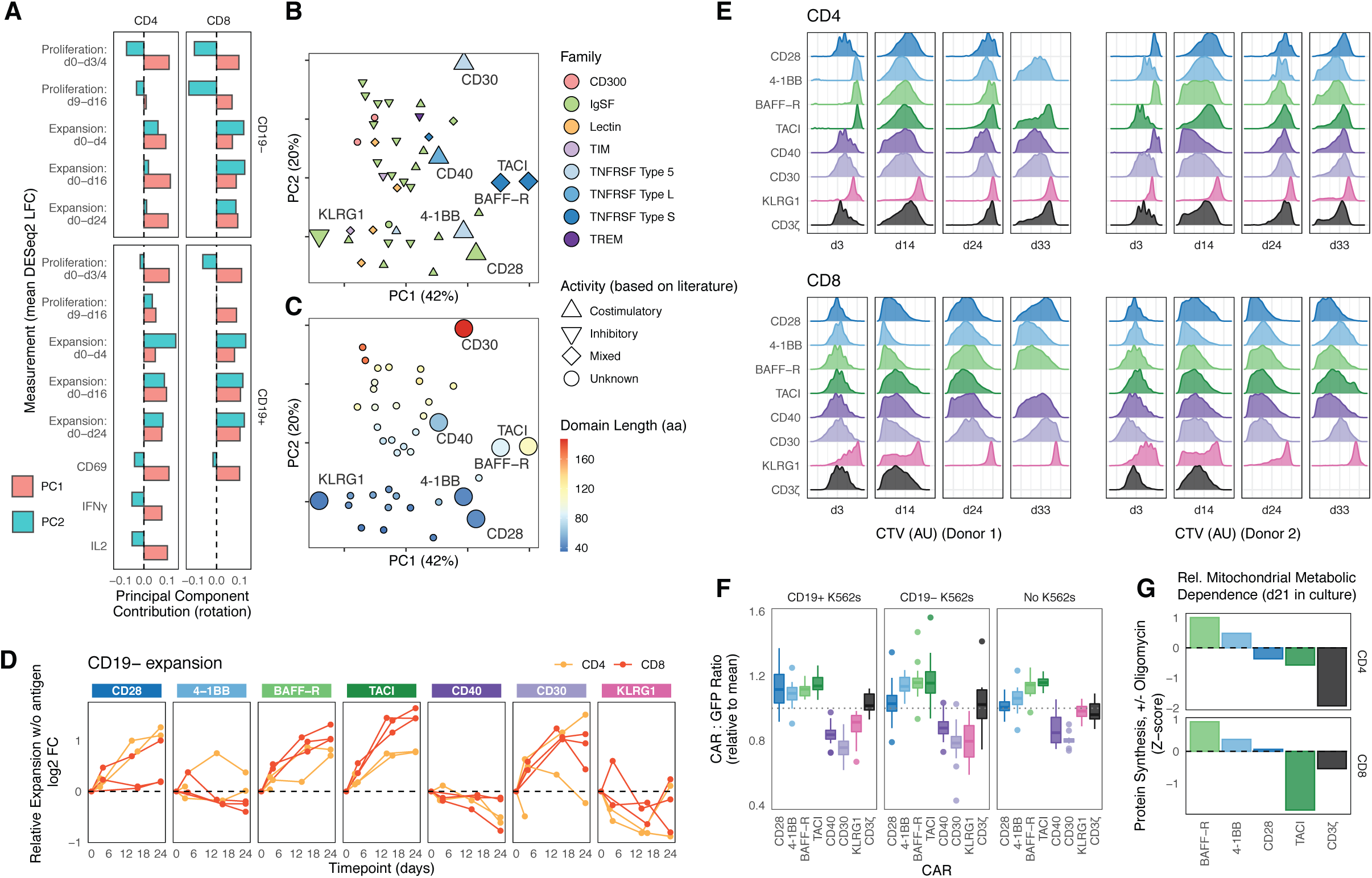
Functional characterization of costimulatory landscape and of proliferation, CAR expression, and metabolism of CARs with chosen costimulatory domains. **A)** A bar plot showing the relative contributions of different measurement types (in CD4s and CD8s, with and without antigenic stimulation) to each principal component of the PCA plot in Figure 3A, S2B, and S2C. Contributions are grouped across donor replicates and separated out by different timepoints, proliferation (CTV FlowSeq), expansion (change in relative library abundance over time), intracellular cytokine FlowSeq, and activation (CD69 FlowSeq). **B)** A recoloring of Figure 3A according to the domain activities (shapes) and domain types (colors) across the library as listed in Figure 1A. Chosen CARs are shown as larger symbols. **C)** A recoloring of Figure 3A according to the amino acid length of each costimulatory domain, showing a correlation between domain length (blue to red is shortest to longest) and the second principal component. **D)** Relative expansion of library members CD28, 4-1BB, BAFF-R, TACI, CD40, CD30, and KLRG1 over 24 days of repeated stimulation with irradiated CD19- K562 cells, as in Figure 3B. Expansion was quantified by calculating the fold-change of the proportion of each CAR within the library at each timepoint (X axis) as compared to baseline relative to the average CAR within the pooled library. The library was measured in CD4 and CD8 primary human T cells individually in 2-3 biological replicates. **E)** Cell Trace Violet flow cytometry histograms, as in Figure 3C, for both donors, all T cell types, and all time points. **F)** Ratio of surface CAR expression (via myc tag Flow Cytometry staining) to GFP fluorescence for each CAR. All CAR variants were normalized to the mean within each time point, donor, and T cell type (CD4 or CD8). Expression with CD19+ K562s, CD19- K562s, and no target cells are shown separately. **G)** Normalized relative metabolic mitochondrial dependence for CD4 and CD8 T cells measured among select CARs. This metric is based on measurement of protein synthesis via the SCENITH method, which calculates the change in overall metabolic output with and without the addition of oligomycin, a mitochondrial inhibitor.

**Figure S3:**
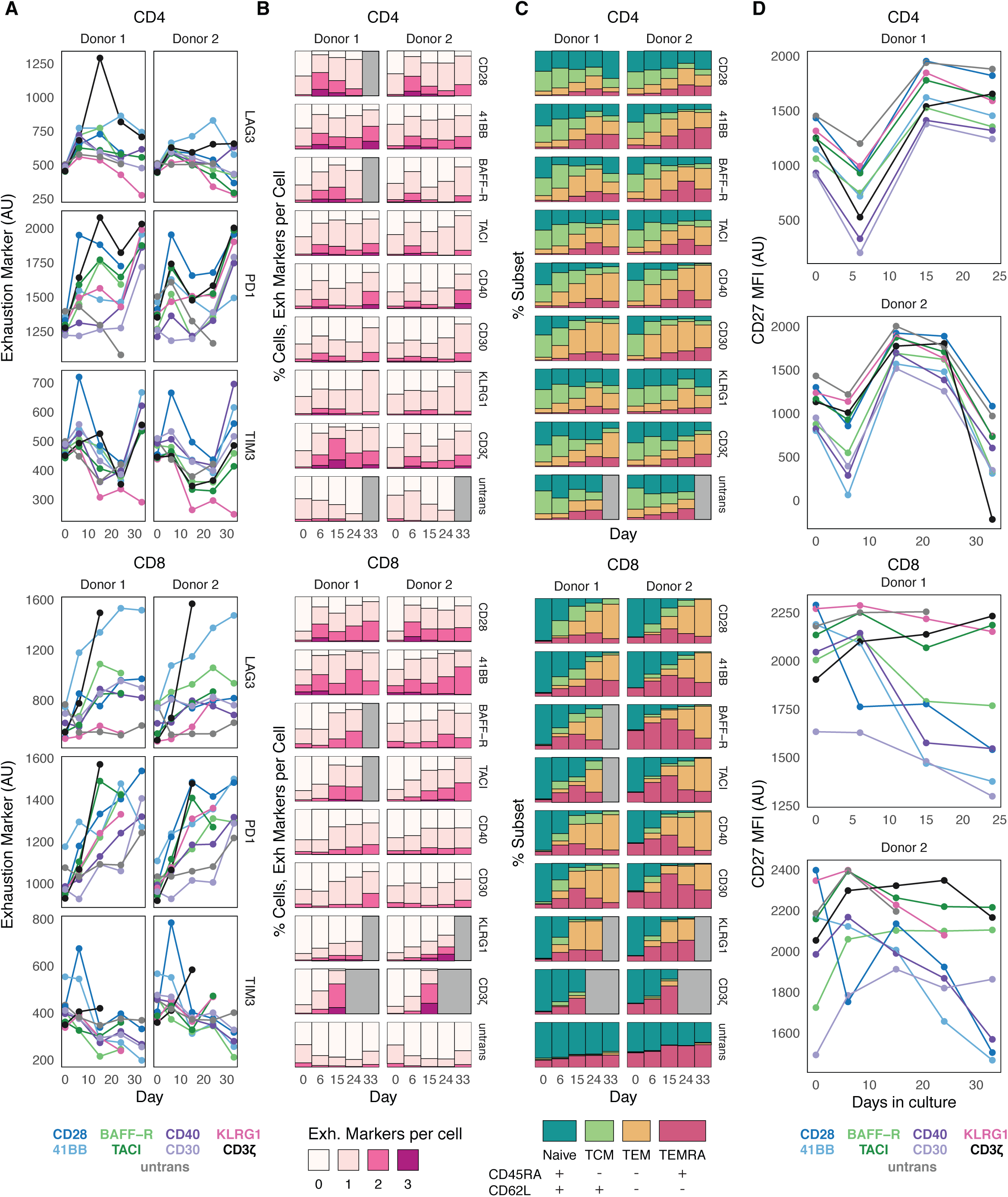
Exhaustion and differentiation characteristics of CARs with chosen costimulatory domains. **A)** Mean fluorescence intensity (MFI) for three cell surface markers of exhaustion (LAG3, PD1, TIM3) in anti-CD19 CAR T cells generated from two donors, measured after repeated stimulation with CD19-expressing irradiated K562s. **B)** Number of CAR T cells expressing 0-3 of the exhaustion markers PD1, TIM3, and LAG3 after different numbers of days in culture, as in Figure 3I, based on the protocol described in Figure 3G. **C)** Differentiation of T cells at different timepoints throughout the repeated stimulation assay. Differentiation subsets (Naive, Central Memory (TCM), Effector Memory (TEM), Effector Memory RA-positive (TEMRA)) were calculated using surface expression of CD45RA and CD62L, as shown in the table at the bottom of the panel. **D)** MFI of CD27 across all T cells, timepoints, and CAR T variants, as in Figure 3I.

**Figure S4:**
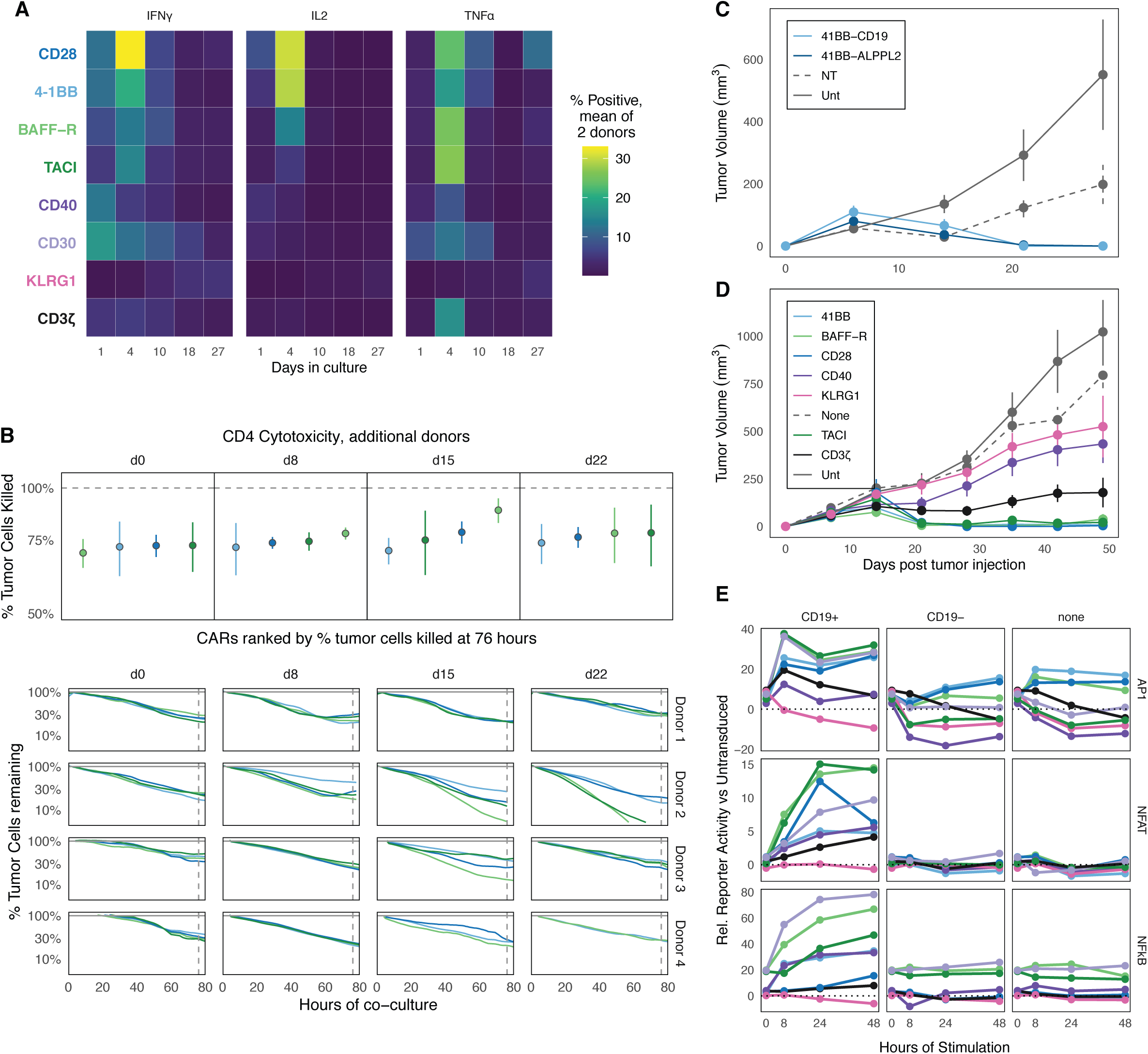
Time course of cytokine production, cytotoxicity, and transcriptional activity across CARs with chosen costimulatory domains. **A)** Mean cytokine production is shown across all T cells, time points, and CAR T variants, as in Figure 4A. **B)** Cytotoxicity of CD4 CAR T cells were quantified at 80 hours (top) with representative plots (bottom) for all four donors expressing BAFF-R, TACI, CD28, or 4-1BB as in Figure 4C and Figure 4D. Colors for each CAR are as labeled in Figure S4A. CARs are ranked at each timepoint from least to most cytotoxic (left to right). Vertical dashed lines indicate the time points analyzed. **C)** We injected 4e6 M28 mesothelioma tumor cells subcutaneously into the flank of NSG mice as in Figure 4E-G and S4D . Seven days later we intravenously injected 6e6 engineered 4-1BB CAR T cells targeting either ALPPL2 or CD19. Untransduced T cells and non-treated mice were included as controls. Tumors were measured via caliper every 7 days for a total of 30 days. **D)** Tumor size is shown over 49 days post tumor injection across all T cells, time points, and TRAC-knockout CAR T variants, as in Figure 4F-4G. **E)** Percent transcription factor activity relative to untransduced reporter Jurkat cells is shown across all time points and CAR T variants as in Figure 4H. Colors for each CAR are as labeled in Figure S4A. Each column indicates a distinct stimulation, either CD19+ K562 cells (left), CD19- K562 cells (middle), or media only (right).

**Figure S5:**
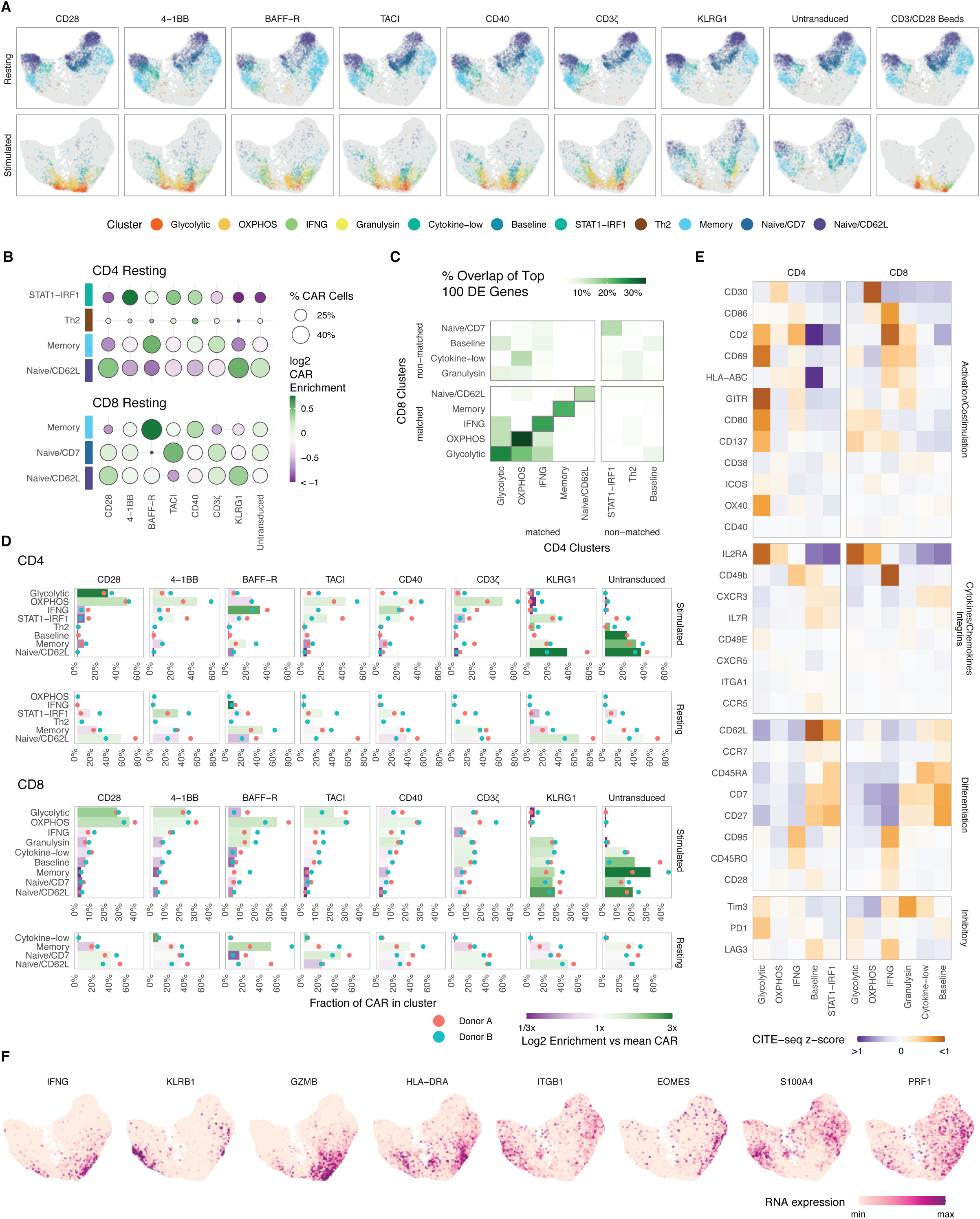
Single-cell analysis of CARs with chosen costimulatory domains with and without antigen stimulation. **A)** UMAP plots faceted separately for each CAR costimulatory domain and stimulation condition. Points are colored the same as Figure 5A. An additional CD3/CD28 bead stimulation condition is also shown, which was done only in Donor 2. **B)** Gene expression overlap across 5 pairs of clusters which are very similar between CD4s and CD8s (Naive/CD62L, Memory, IFNG, OXPHOS, and Glycolytic). A list of the top 100 differentially expressed genes was calculated for each cluster among all CD4 or CD8 cells. This plot shows the percentage overlap in these gene lists between clusters, showing a marked mirroring of gene expression across the CD4/CD8 axis among the 5 matched clusters in the bottom left quadrant. **C)** Enrichment of resting CAR T cells containing different cosignaling domains within each phenotypic cluster, similar to Figure 5C. The size of each dot corresponds to the percentage of stimulated CAR T cells with a specific costimulatory domain that is assigned to a cluster. The color of each dot corresponds to the log-2 fold enrichment or depletion of that CAR within the cluster. **D)** CITE-seq z-scores for a variety of surface proteins among T cells in different activated clusters, grouped by their functional classification. Z-scores for CD4 and CD8 T cells were calculated separately. **E)** A breakdown of the cluster frequency among all stimulated and resting CAR variants of both donors. The bar length on the X axis is the percentage of each costimulatory CAR variant (resting and stimulated separately) within that cluster, such that each set of bars within each faceted box sums to 1. The bars represent the mean percentage for both donors, while the blue and red dots represent the individual percentages for each donor. The color of each bar corresponds to the relative log2 enrichment for that CAR variant in that cluster, relative to other CAR variants. **F)** UMAP heatmaps display the relative RNA expression of single cells (scaled individually), showing a subset of functionally-important transcripts that are upregulated in the IFNG and Memory subsets.

**Supplemental Table 1.**
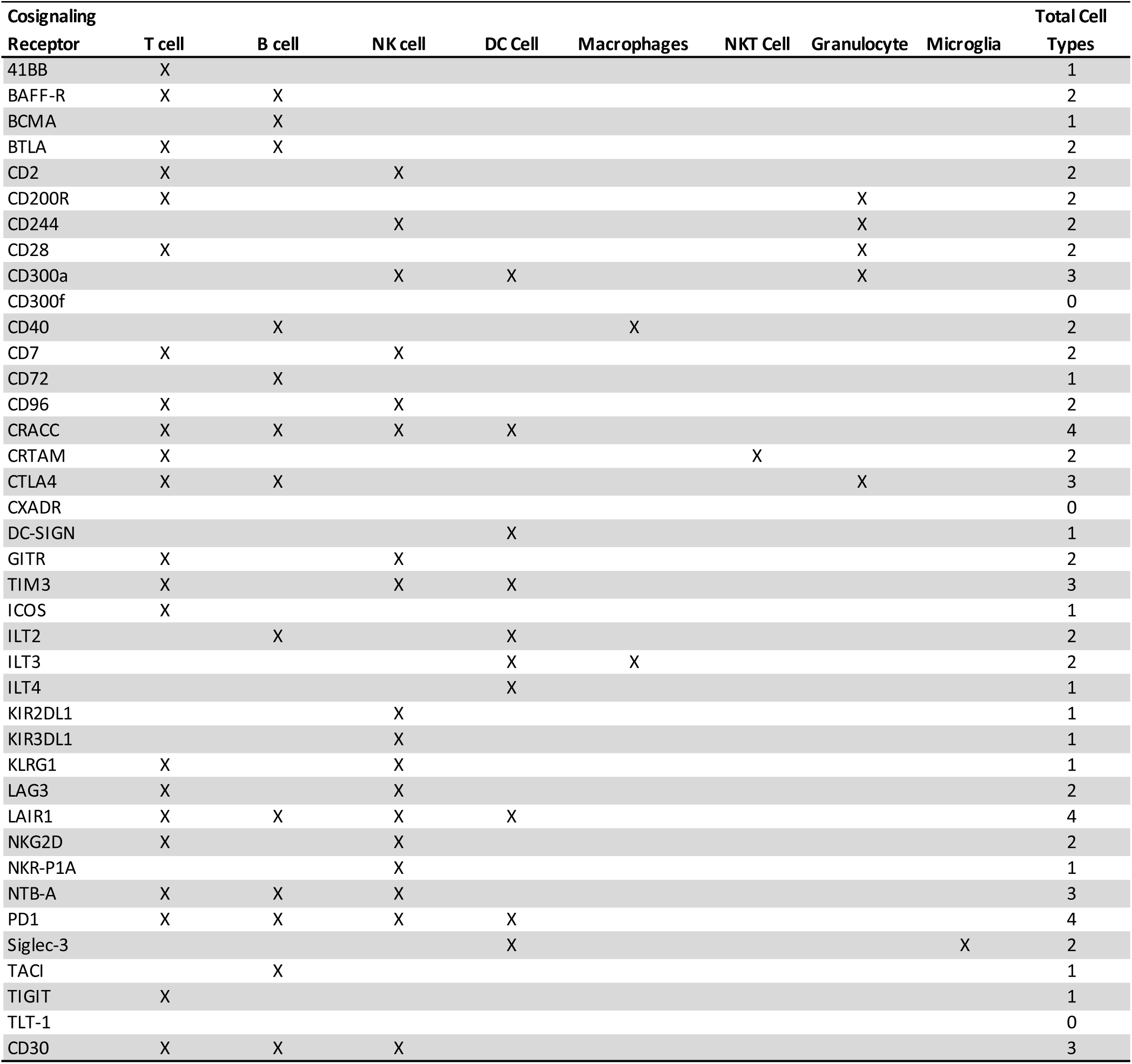
Expression of individual cosignaling receptors by cell type. A list of all costimulatory domains in our library and whether they are expressed by different immune cell types. Note that some receptors may have low expression or only be expressed under specific circumstances by individual cell types.

